# Generation of cardiac valve endocardial like cells from human pluripotent stem cells

**DOI:** 10.1101/2020.04.20.050161

**Authors:** LX Cheng, Y Song, YY Zhang, YC Geng, WL Xu, Z Wang, L Wang, K Huang, NG Dong, YH Sun

## Abstract

The cardiac valvular endothelial cells (VECs) are an ideal cell source that could be used for the fabrication of the next generation tissue-engineered cardiac valves (TEVs). However, few studies have been focused on the derivation of this important cell type. Here we describe a chemically defined xeno-free method for generating VEC-like cells from human pluripotent stem cells (hPSCs) through an intermediate endocardium stage. HPSCs were initially specified to KDR^+^/ISL1^+^ multipotent cardiac progenitor cells (CPCs), followed by differentiation into endocardial progenitors under the combined treatment with VEGFA, BMP4 and bFGF. In the presence of VEGFA, BMP4 and TGFb, valve endocardial progenitor cells (VEPs) were efficiently derived from endocardial progenitors without a sorting step. Mechanistically, administration of TGFb and BMP4 may specify the VEP fate by facilitating the expression of key transcription factors ETV2 and NFATc1 at the immediate early stage and by activating Notch signaling at the later stage. Notch activation is likely an important part of VEP induction. HPSC-derived VEPs exhibited morphological, molecular and functional similarities to that of the primary VECs isolated from normal human aortic valves. When hPSC-derived VEPs were seeded onto the surface of the de-cellularized porcine aortic valve (DCV) matrix scaffolds, they exhibited higher proliferation and survival potential than the primary VECs. Our results suggest that hPSC-derived VEPs could serve as as a potential platform for the study of valve development, and as starting materials for the construction of the next generation TEVs.

**Highlights:**

1. Valve endocardial progenitor cells (VEPs) could be efficiently derived from hPSCs without a sorting step
2. The combined treatment with TGFb and BMP4 induce VEP fate by enhancing the expression of *ETV2* and *NFATc1*
3. HPSC-derived VEPs resemble the isolated primary VECs molecularly, morphologically and functionally
4. HPSC-derived VEPs exhibit proliferative and functional potential similar to the primary VECs when seeded onto the DCVs

## Introduction

Valve disease is one of the most common cardiac defects, characterized by inflammation, sclerotic lesion, calcification, myxomatous and thrombus formation (Combs and Yutzey, 2009). There are currently no good medical treatments for dysfunctional valves, leaving surgical replacement with mechanical and bioprosthetic valves as the major option. However, the current mechanical and bioprosthetic valves have their limitations due to the requirement of lifelong anti-coagulation, poor durability, and lack of self-growth and self-renewal capacity (Hu et al., 2013). The next generation tissue engineered heart valves (TEVs) which offer a potential remedy to the challenge are highly demanded (Zhou et al., 2013).

Fabrication of the next generation TEVs requires the generation of sufficient number of valvular-like cells (Glaser et al., 2011; Nejad et al., 2016). Cardiac valve is composed of two major cell types: the valvular endothelial cells (VECs) lining the outer surface of the valve cusps and the valvular interstitial cells (VICs) embedded in a stratified extracellular matrix. HPSCs are known to have the potential for unlimited expansion and differentiation into any cell types in a petri dish. A growing body of research has shown that cardiovascular endothelial cells could be efficiently generated from hPSCs (Wilson et al, 2014). Surprisingly, few studies have been focused on the derivation of valve-specific endothelial cells: the VECs (Neri et al., 2019). Thus, it is of great importance in the field to establish a hPSC based in vitro differentiation model that recapitulates VEC lineage fate determination, which would facilitate efficient and reproducible generation of VEC-like cells for potential applications in valve disease modeling, drug screening, and construction of the next generation TEVs.

From the viewpoint of developmental biology, a reliable approach to generate the VEC-like cells would be to recapitulate the embryonic valvulogenesis under chemical defined medium based on the comprehensive understanding of the signaling intricacies involved in the process. Genetic studies using mice and zebrafish models have suggested that the majority of valvular cells are of endocardial cell origin (Hinton and Yutzey, 2011). The endocardial cells (ECs) are a spatially restricted cell population by de novo vasculogenesis from cardiogenic progenitors distinct from the hemangioblasts that give rise to other endothelial subtypes. A growing body of studies support the notion that the endocardial cells are derived from *Isl1^+^*/*Kdr^+^* multipotent progenitors via cardiogenic mesoderm cells expressing *T^+^*/*Mesp1^+^* (Andersen et al., 2018; Nakano et al., 2016; Hoffman et al., 2011).

At E9.5 of mouse embryos or day 30 of human fetus, a subset of endocardial cells at defined regions of the atrio-ventricular (AV) canal and the outflow tract (OFT) (referred as valve endocardial progenitor cells/VEPs) are induced to loose the contact with neighboring cells and undergo an EndoMT to form the heart valve mesenchymal cells (Wu et al., 2011). Although little is known about the valve endocardial cells, studies from mice and zebrafish have shown that they may bear a unique gene expression profile, being expressing the valve endocardial marker genes (VEGs) such as *NAFTc1*, *TBX2*, *TBX3*, *ENG*, *EMCN*, *MSX1*, *SELP*, *HAPLN1*, *TGFβ1/2*, *NOTCH1/3*, *HEY1/2* and *JAG1* (Holliday et al., 2011; Hulin et al., 2019; Lincoln et al., 2006; Neri et al., 2019; Wu et al., 2011). Previous studies have shown that components of multiple signaling pathways (such as BMP, FGF, Notch and TGF-β) and transcription factor such as ETV2/NfATc1/GATA4 are expressed in a spatial-temporal manner during endocardial cushion formation, the EndoMT and valve remodeling (Garside et al., 2013; Wu et al., 2011; Yang et al., 2017). For instance, BMP ligands 2, 4, 5, 6, and 7 as well as BMP type I receptors are all expressed in the AVC and OFT prior to valve formation (Jiao et al., 2003; Kruithof et al. 2012; Liu et al., 2004; Nakano et al, 2012; Stainier et al., 2002). BMP activation promotes the cardiac differentiation of endothelial progenitors (Sahara et al., 2014; Sriram et al., 2015), while attenuating Bmp signaling inhibited SHF differentiation and led to cushion defects (Brade et al., 2013; Nakano et al., 2016; Kattman, 2011; Rochais et al., 2009; Palpant et al., 2015; Combs and Yutzey, 2016; Hinton and Yutzey, 2011). The requirement for FGF signaling in endocardial cushion morphogenesis and endocardial cell derivation have been reported (Neri et al., 2019; Park et al., 2008), and FGF deficient embryos have lethal malformations of the cardiac OFT and heart (Frank et al., 2002; Marques et al., 2008). TGF ligands and receptors are expressed in the AVC and OFT during endocardial cushion formation in avian and murine embryos (Lamouille et al., 2014). TGFb-null embryos demonstrate severe cardiac abnormalities, including disorganized valves and vascular malformations (Hinton and Yutzey, 2011; Laforest and Nemer, 2012). Although signaling pathways and transcription factors between the endocardium and the myocardium play important roles in the formation of the valve endocardial cells, the mechanisms by which the VEP fate is specified and determined remain largely unclear (Hinton and Yutzey, 2011;McCulley et al., 2008; Wu et al., 2011, 2012).

In this work, we reported that using a chemically defined xeno-free method, VEC-like cells can be efficiently derived by differentiating hPSCs (H8, H9 and iPSCs) into an endocardial cell fate. HPSC-derived VEC-like cells exhibited morphological, molecular and functional similarities to that of the primary VECs isolated from normal human aortic valves. When hPSC-derived VEC-like cells were seeded onto the surface of the de-cellularized porcine aortic valve (DCV) matrix scaffolds, they exhibited proliferation and survival potential similar to the primary VECs. Our results suggest that hPSC-derived VEPs could serve as as a potential platform for the study of valve development, and as starting materials for the construction of the next generation TEVs.

## Results

### Efficient generation of ISL1^+^ cardiac progenitor cells from hPSCs

It is widely accepted that the mammalian embryonic heart is primarily originated from cardiac progenitor cells (CPCs) expressing marker genes such as *ISL1*, *NKX2.5* and *KDR* (Hinton and Yutzey, 2011; Lui et al., 2013; Misfeldt et al., 2009; Morreti et al., 2006). We and several other groups have previously generated the CPCs from hPSCs (Berger et al., 2016; Colunga et al., 2019; Lyer et al., 2015; Nakano et al., 2016; Patsch et al., 2016; Sahara et al., 2014; Sriram et al., 2015; Tan et al., 2013; White et al., 2013). The common theme of the various differentiation protocols is the activation of WNT and BMP signaling pathways which mimics the signaling requirement during early specification of heart mesoderm (Ansersen et al., 2018; Nakano et al., 2016; Puceat, 2013).

It has been shown that there is a greater propensity for ISL1^+^ cardiac progenitors to generate endocardium when NKX2.5 was low or absent (Morreti et al., 2006; Nakano et al., 2016). We therefore aimed to generate ISL1^high^NKX2.5^low^ CPCs from hPSCs. To this end, we have modified our previously published protocol (Berger et al., 2016). HPSCs were initially treated with BMP4 and WNT agonists (WNT3a or the small molecule CHIR99021) for 3 days, and the expression of *T*, a primitive streak and early mesodermal marker, was monitored daily (Figure 1A; Supplementary Figure 1A-B). CHIR99021 is a highly specific glycogen synthase kinase-3 (GSK3) inhibitor that can activate canonical Wnt signaling (Chappell et al., 2013). Compared to the treatment with BMP4 or WNT3a only, combined treatment with BMP4 and WNT3a resulted in a quick induction of *T*, reaching its maximum expression levels at day 1 (Supplementary Figure 1A). Next, day 1 *T^+^* cells were treated with bFGF and BMP4 for 6 days, and qRT-PCR analysis was performed to examine the expression of CPC markers such as ISL1 and NKX2.5. The qRT-PCR results showed that *ISL1* and *KDR* were highly induced, peaking at day 2 post-treatment; while the first heart field (FHF) progenitor markers such as *NKX2.5* and *TBX5* were barely up-regulated (Supplementary Figure 1B,C). Immunofluorescence (IF) staining of day 3 CPCs showed that a vast majority (> 90%) of cells were ISL1 positive while cells displayed negligible expression of NKX2.5 (Figure 1C-D; Supplementary Figure 1D). Western blot analysis confirmed that ISL1 and KDR but not NKX2.5 were abundantly expressed in day 3 hPSC-derived CPCs (Figure 1E). Finally, the flow cytometry analysis further revealed that approximately 80% of day 3 CPCs were KDR positive and over 93% of cells were ISL1 positive, indicating that hPSC cardiogenic mesodermal differentiation was of high efficiency (Figure 1F).

**Figure 1.**
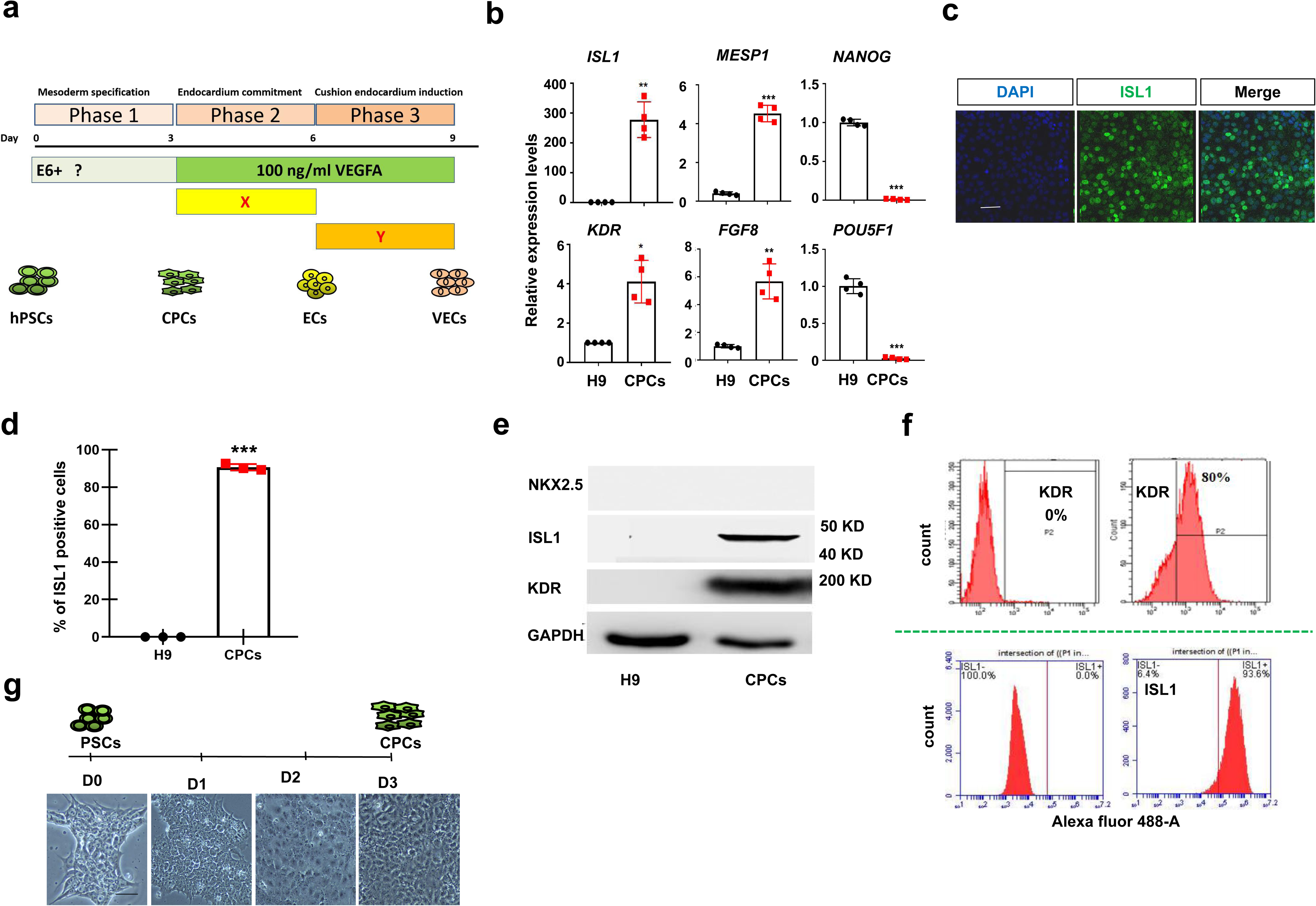
PSCs to cardiogenic mesoderm expressing KDR and ISL1 a. Schematic of differentiating hPSCs to CPCs, endocardial progenitor cells and VEPs. **b** The qPCR analysis of day 3 hPSC-derived CPCs for the indicated markers. **c** IF staining of day 3 CPCs showing that the majority of cells were ISL1 positive. Scale bar: 50 μm. **d** Quantification analysis of panel (c) by ImageJ. Bar graph represents percentage of ISL1 positive cells ± S.D of three independent experiments. **e** WB analysis of day 3 CPCs using the indicated antibodies. **f** Flow cytometry analysis showing percentage of KDR and ISL1 positive cells. **g** The representative morphology of cells daily during hPSC differentiation to CPCs. All experiments were repeated 3 times. Significant levels are: \**p*< 0.05; \*\**P*< 0.01; \*\*\**P*< 0.001. Shown are representative data.

Based on the above results, we concluded that ISL1^high^NKX2.5^low^ CPCs could be efficiently generated from hPSCs in 3 days (Figure 1G).

### Differentiation of CPCs into endocardial cell types

Although the endocardium within the heart is contiguous with the rest of the vasculatures, it is a unique and molecularly and functionally distinct population different from other endothelial subtypes, expressing higher levels of marker genes such as *NFATC1, TEK, GATA4, ETV2, MEF2C, TBX20* and *SMAD6* (Harris and Black, 2010; Neri et al., 2019; Wilson et al., 2012; Puceat, 2013). Studies from several genetic models have suggested that the endocardium arises from KDR^+^/ISL1^+^ multipotent CPCs distinct from the hemangioblasts that give rise to extracardiac endothelial cell types (Misfeldt et al., 2009; Milgrom-Hoffman et al., 2011).

Day 3 hPSC-derived CPCs were initially treated with 100 ng/ml VEGFA for 3 days, and the expression of endocardial markers was monitored by qRT-PCR (Figure 2A). VEGFA is known to be the key signaling factor for PSC differentiation into cardiovascular endothelial cells (Sahara et al., 2014; Sriam et al., 2015; Neri et al., 2019). The qRT-PCR results showed that there was a significant up-regulation of general endothelial cell markers such as *PECAM1* and *CDH5* (Supplementary Figure 2A). The day 3 post-treatment cells also abundantly expressed *TEK, GATA4* and *KDR*, indicating that treatment with VEGFA is sufficient to convert CPCs to cardiovascular endothelial progenitors. Unfortunately, the endocardium-enriched marker genes such as *NFATc1, SMAD6, ETV2* and *TBX20* were barely induced (Figure 2B; Supplementary Figure 2A), which suggested that additional factors are required for the induction of endocardial cell types.

**Figure 2.**
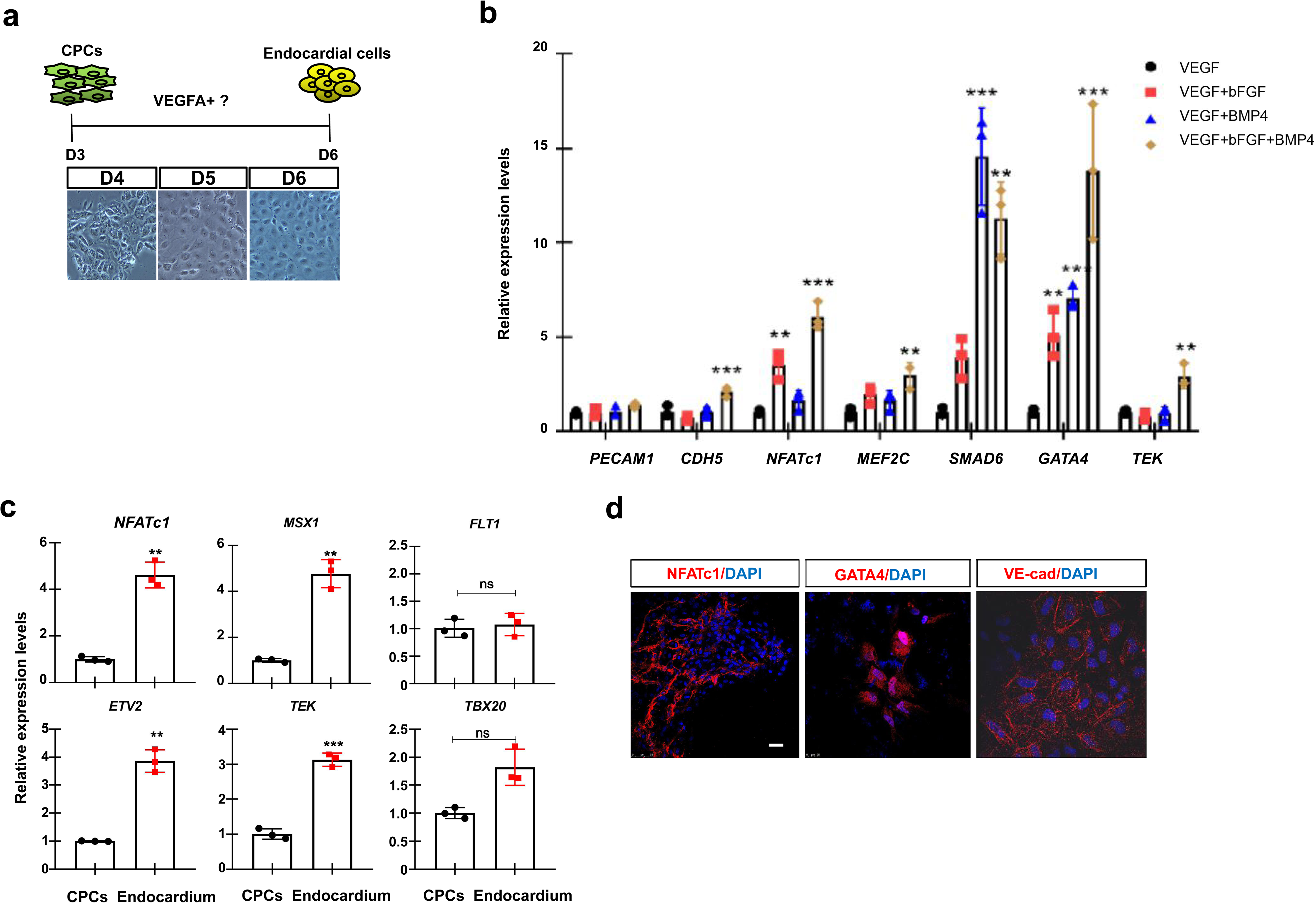
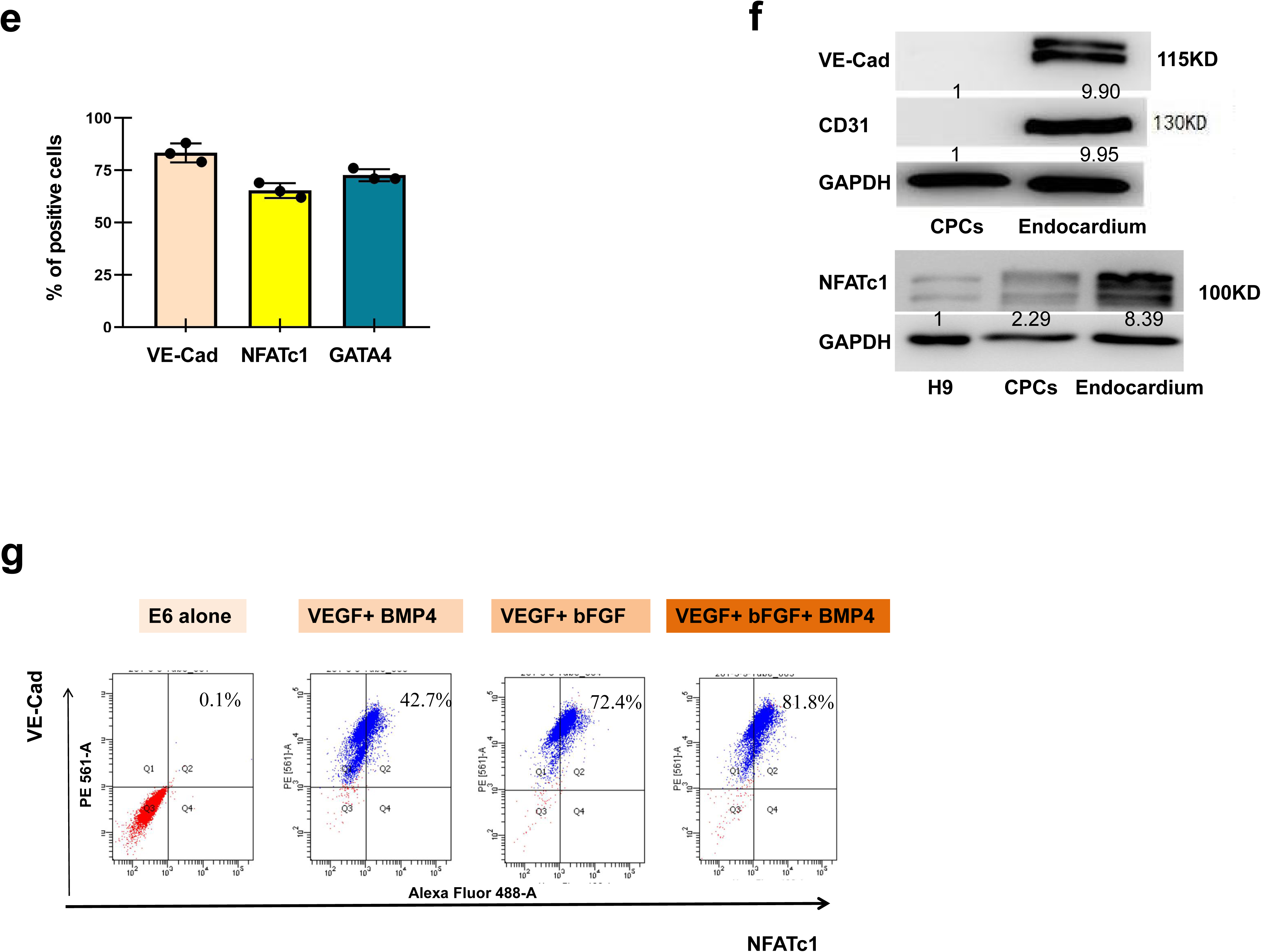
Cardiogenic mesoderm to endocardial endothelial cells a. Schematic of hPSC-derived CPCs differentiation to endocardium progenitor cells. **b** The qPCR analysis of the known endocardium markers for hPSC-derived CPCs that were treated with the indicated signaling molecules for 3 days. **c** The qRT-PCR analysis of the known endocardium markers of hPSC-derived CPCs that were treated with VEGFA, BMP4 and bFGF for 3 days. **d** IF staining of day 6 hPSC-derived endocardial progenitors showing abundant expression of VE-Cad, GATA4 and NFATc1. Scale bar: 25 μm. **e** Quantification analysis of panel (d) by ImageJ. Bar graph represents percentage of VE-Cad, GATA4 and NFATc1 positive cells ± S.D of three independent experiments. **f** Western blotting using the indicated antibodies showing strong expression of VE-Cad, CD31 and NFATc1 in day 6 hPSC-derived endocardial progenitors. **g** Flow cytometry analysis showing percentage of VE-Cad and NFATc1 double positive cells. Day 3 hPSC-derived CPCs were treated with the indicated signaling molecules for 3 days, and stained with VE-Cad and NFATc1 antibodies. All experiments were repeated 3 times. Significant levels are: \**p*< 0.05; \*\**P*< 0.01; \*\*\**P*< 0.001. Shown are representative data.

To identify the signaling conditions that promote CPC to an endocardial lineage, we systematically analyzed the effects of various signaling molecules and modulators (including the activators and inhibitors of TGF-β/Activin A, FGF, Notch, WNT and BMP signaling pathways). HPSC-derived CPCs were trypsinized and replated as a monolayer for 3 days under the treatment with low and high concentrations of different growth factors or modulators, and the expression of endocardium-enriched markers was examined (Supplementary Figure 2A). Remarkably, treatment with bFGF led to significantly increased expression of *CDH5* and *PECAM1* as well as endocardial marker genes such as *NFATc1*, *TBX20* and *MEF2C*, without obvious effect on the expression of fibroblast markers such as *POSTN, ACTA2/SMA* and *S100A4*/*FSP1*. The results were in line with that VEGF and bFGF are two critical cytokines for cardiogenic mesoderm commitment to an EC fate (Lui et al., 2013; Neri et al., 2019; Nolan et al., 2013; White et al., 2013), and that FGFs are required to generate the pre-valvular endocardial cells from hPSCs (Neri et al., 2019). BMP4 has been shown to be required for cardiovascular endothelial/endocardial differentiation (Palencia-Desai et al., 2015; Sahara et al., 2014), and enhance hPSC differentiation into ECs in a dose dependent fashion (Sriram et al, 2015; White et al., 2013). We found that combined treatment with VEGFA and BMP promoted the expression of endocardial markers *NFATc1, TBX20* and *MEF2C,* with little effect on the fibroblast markers *POSTN* and *S100A4*/*FSP1* (Supplementary Figure 2A). Administration of TGFb/Activin-A barely induced *NFATc1* and *MEF2C* but led to a dramatic up-regulation of cardiac fibroblast markers *VIM* and *POSTN.* Addition of WNT3A or CHIR failed to up-regulate the endocardial markers but had a significant role in promoting *POSTN* and *VIM* expression, which was in line with previous reports showing that WNT activities need to be restrained for an EC lineage (Palpant et al., 2015; Nakano et al., 2013) (Supplementary Figure 2A, B). Notch signaling has been shown to play a key role in ESC differentiation towards an EC fate (Sahara et al., 2014; Wang et al., 2013). DLL4 is a transmembrane ligand for Notch receptors, and the γ-secretase inhibitor DAPT is a potent Notch signaling inhibitor (Pompa et al., 2009). We surprisingly found that treatment with either DLL4 or DAPT failed to have a significant effect on the expression of endocardial markers except for *NFATc1* (Supplementary Figure 2B, C). Based on the pilot experiments, we primarily focused on the effects of modulating BMP and FGF signaling pathways in the subsequent experiments.

Next, hPSC-derived CPCs were treated with 100 ng/ml VEGFA along with increasing dose (10, 20, 50 and 100 ng/ml) of bFGF, and the dynamic expression of endocardial markers was monitored over a time course of 12 days. Of all the doses tested, 20 ng/ml bFGF constantly produced the best results in our hand, as revealed by mRNA expression analysis of the selected endocardial markers (Supplementary Figure 2D). In the combined treatment with VEGFA and bFGF, the expression of definitive endothelial and endocardial maker genes was gradually increased, peaking at day 6 (Figure 2B; Supplementary 2E). Consistently, these cells adopted and remained an EC like cobblestone morphology after 3 days of the treatment (Supplementary Figure 2F). When the FGF/MEK inhibitor PD0325901 (PD) or U0126 (U0) was applied, the expression of endocardial genes was significantly decreased (Supplementary Figure 2G). To determine the optimal BMP4 dose, hPSC-derived CPCs were treated with varying concentrations (5, 10, 25, 50 and 100 ng/ml) of BMP4 in the presence of VEGFA. In general, BMP4 treatment up-regulated the endocardial marker genes in a dose dependent manner. However, higher concentration of BMP4 also induced the fibroblast markers *POSTN*, *BGN* and *ACTA2* (Supplementary Figure 2A, H). We thus determined 5-10 ng/ml BMP4 as the optional concentration as it promoted the expression of endocardial markers while failed to induce fibroblast markers. Actually, this dose of BMP4 was very close to that in a previous study (White et al., 2013). When BMP signaling inhibitor LDN193189 (LDN) was applied, the mRNA levels of endocardial marker genes were significantly decreased (Supplementary Figure 2I).

Next, we investigated the effect of the combined treatment with VEGFA, BMP4 and bFGF on day 3 hPSC-derived CPCs for 12 days. The treated cells adopted and remained an EC like morphology over days 3-12 (data not shown). When day 6 cell cultures were collected and used for gene expression analysis, combined treatment with bFGF and BMP4 led to much higher expression levels of endocardial markers compared to the treatment with bFGF or BMP4 only (*P*< 0.05), suggesting that there was an additive action between bFGF and BMP4 on the endocardial markers (Figure 2B, C). Importantly, these cells exhibited minimal expression levels of cardiac fibroblast markers *POSTN* and *ACTA2*, cardiomyocyte marker *TNNT1* as well as epicardial markers *WT1* and *TBX18* (Supplementary Figure 2j). In addition, human hematopoietic stem cell markers *CD133*/*PROM1* and *CD45*/*PTPRC* were not induced, suggesting that hPSC-derived CPCs were primarily differentiated towards an endocardial but not hemogenic EC lineage (Supplementary Figure 2K). IF staining of day 6 hPSC-derived endocardial-like cells showed that they abundantly expressed GATA4, VE-cad and NFATc1 (Figure 2D). Quantitative analysis of the immunofluorescence (IF) staining showed that approximately 85% of the cells were VE-cad positive, 81% were GATA4 positive, and 80% were NFATc1 positive (Figure 2E). WB analyses confirmed that day 6 hPSC-derived endocardial-like cells expressed GATA4, VE-cad and NFATc1 at protein levels (Figure 2F). Finally, the flow cytometry results revealed that approximately 82% of day 6 hPSC-derived endocardial cells were NFATc1 and VE-cad double positive (Figure 2G), which indicated that hPSC endocardial cell differentiation was of high efficiency. Note that the percentage of NFATc1 and VE-cad double positive cells was significantly higher under the combined treatment with bFGF and BMP4 than treatment with bFGF or BMP4 only.

Taken together, our results demonstrated that endocardial cell types can be efficiently obtained when hPSC-derived CPCs are differentiated under the combined presence of VEGFA, BMP4 and bFGF for three days.

### Specification of endocardial cells to valve endocardial progenitors

Following the formation of the endocardial layer, the cardiac tube forms and undergoes looping morphogenesis. Soon after looping, endocardial cells at defined regions of the AVC and the OFT lose the contact with neighboring cells. It is these AVC/OFT endocardial populations (valve endocardial progenitors/VEPs) that undergo an EndoMT process to form the interstitial cells of the future valve cusps. Lineage tracing studies in several models have shown that components of multiple signaling pathways (such as ligands and receptors of BMP, Notch, FGF and TGF-β signaling pathways) are expressed around the endocardial cushion or its environment, which strongly suggests that they play important roles for VEP specification and fate decision (Hinton and Yutzey, 2013; Latif et al., 2015; McCulley et al., 2008; Pompa et al., 2009). We therefore investigated the effects of modulating BMP, FGF, TGF-β and Notch signaling on day 6 hPSC-derived endocardial progenitors (Figure 3A).

**Figure 3.**
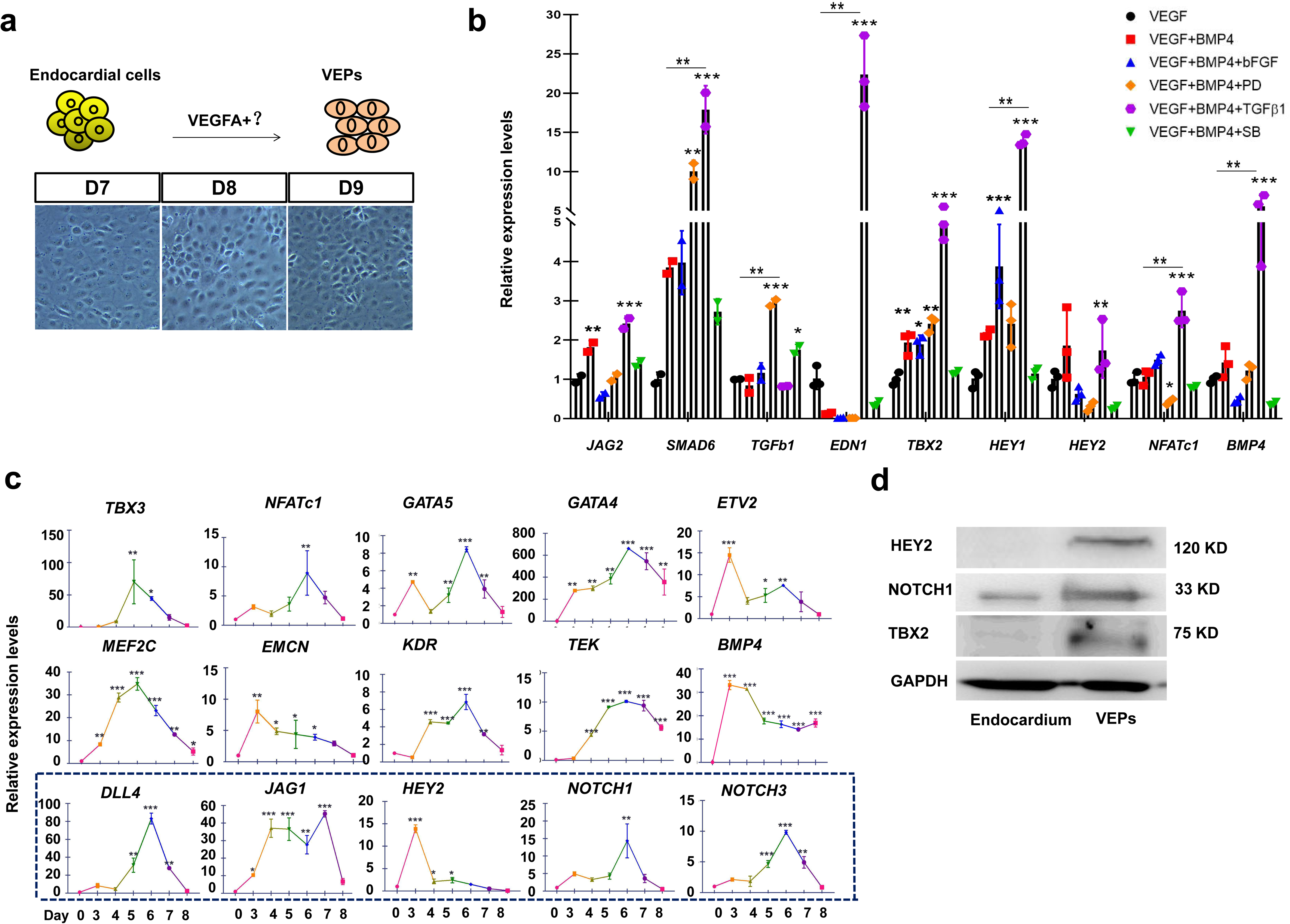
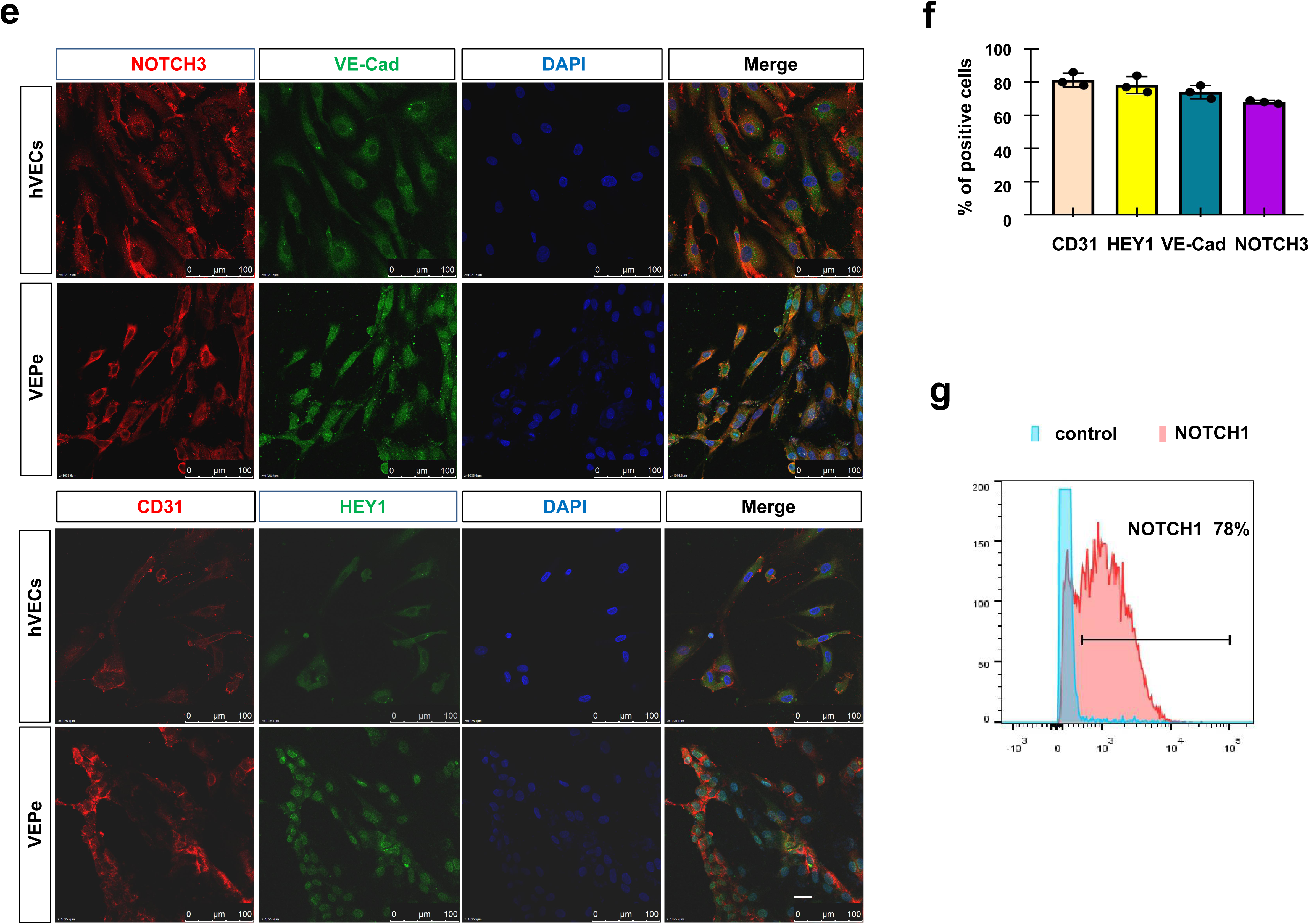
Endocardium progenitors to AVC endocardial cells a. Schematic of hPSC-derived endocardial cells specification to the VEPs. **b** The qPCR analysis of the selected VEGs for day 9 hPSC-derived VEPs. Note that the combined treatment with VEGF, BMP4 and TGFb leads to a dramatic up-regulation of the VEGs. **c** The dynamic expression analysis of known VEGs of hPSC-derived endocardium cells that are treated with BMP4 and TGFb for a time course of 9 days. The Notch-related markers were high-lighted by a purple dashed box. **d** WB analysis for hPSC-derived endocardial cells and VEPs, using the indicated antibodies. **e** IF staining of day 9 hPSC-derived VEPs and the primary VECs, showing that a portion of the cells are positive for VE-cad, CD31, HEY1 and NOTCH3. Scale bar: 25 μm. **f** Quantification analysis of panel (e) by ImageJ. Bar graph represents percentage of VE-cad, CD31, HEY1 and NOTCH3 positive cells ± S.D of three independent experiments. **g** Flow cytometry result showing the percentage of NOTCH1 positive cells. All experiments were repeated 3 times. Significant levels are: \**p*< 0.05; \*\**P*< 0.01; \*\*\**P*< 0.001. Shown are representative data.

Initially, day 6 hPSC-derived endocardial progenitors were treated with BMP, FGF, TGF-β and Notch agonists or antagonists for 3 days in the presence of VEGFA, and qRT-PCR analyses were performed to monitor the expression of selected VEGs such as *NAFTc1*, *TBX2*, *TBX3*, *TGFβ1*, *HEY1/2* and *JAG1/2* (Holliday et al., 2011; Hulin et al., 2019; Lincoln et al., 2006; Neri et al., 2019) (Figure 3B; Supplementary Figure 3A). Although Notch signaling has been shown to be implicated in the endocardial cushion induction and valve cusp formation, modulating Notch signaling (by the addition of different doses of the Notch ligand DLL4 or the Notch signaling inhibitor DAPT) had a minimal effect on the expression of the VEGs (Supplementary Figure 3A). Similar results were observed when FGF/ERK signaling was modulated by treatment with different doses of bFGF or FGF/MEK inhibitor PD (Figure 3B; Supplementary Figure 3A). It has been shown that an increased levels of both TGFb and BMP4 occur in the endocardial cells overlying the cushions of the AV canal and OFT (Kruithof et al., 2012; McCulley et al., 2008). In addition, the combined treatment with BMP4 and TGFb was shown to promote hPSC differentiation into cardiovascular endothelial cell types (Di Bernardini et al., 2014; White et al., 2013). Thus, previous studies strongly suggest that a crosstalk of BMP4 and TGF signaling might be required for the VEP fate specification. We tested this possibility and found that this was the case. Although administration of BMP4 or TGFb had a minimal effects on mRNA expression of the selected VEGs, combined treatment with BMP4 and TGFb led to a marked up-regulation of these marker genes (Figure 3B; Supplementary Figure 3A, B). Thus, our preliminary data suggested that a crosstalk (synergy) between the BMP and TGF-β signaling pathways might be involved in the specification of valve endocardial-like cell types (Hinton and Yutzey, 2013; Nolan et al., 2013).

To gain more insights into the effects of combined treatment with BMP4 and TGFb, the dynamic expression of VEGs was carefully monitored by qRT-PCR over a time course of 9 days. As shown in Figure 3C, the expression of VEGs was gradually increased, peaking at day 3-6 post-treatment. Interestingly, Notch-related genes such as *NOTCH1/3*, *JAG1*, *DLL4* and *HEY2* and its target genes such as *HAND2/TBX2/3* were all significantly induced (VanDusen et al, 2014) (Figure 3C, D; Supplementary Figure 3A, B), suggesting that Notch signaling is activated following the combined treatment with BMP4 and TGFb. The treated cells adopted and remained an EC-like cobblestone morphology (Supplementary Figure 3C). As day 9 hPSC-derived cell cultures (day 3 post-treatment of day 6 hPSC-derived endocardial cells) exhibited the maximum expression levels of VEGs, they were chosen for further analyses as described below. WB analyses showed that valve endocardial markers such as HEY2, NOTCH1 and TBX2 were more strongly induced in day 9 cell cultures compared to day 6 hPSC-derived endocardial cells (Figure 3D). IF staining results showed that day 9 hPSC-derived cells abundantly expressed CD31, VE-cad, NFATc1, NOTCH3 and HEY1, similar to the primary VECs freshly isolated from normal human aortic valves (Figure 3E). Quantitative analysis of IF staining revealed that approximately 70-80% of the cells were NFATc1, NOTCH3, HEY1, VE-cad and CD31 positive (Figure 3F; Supplementary Figure 3D-F). Finally, flow cytometry analysis of day 9 hPSC-derived cells revealed that approximately 78% of the cells were NOTCH1 positive (Figure 3G). Thus, day 9 hPSC-derived cells displayed a gene expression profile similar to the expression signature of the pre-valvular endocardium (Hulin et al., 2019; Neri et al., 2019). Based on the above results, we tentatively named day 9-10 hPSC-derived cells as valve endocardial progenitors (VEPs).

Taken together, our data strongly suggested that under combined treatment with VEGFA, BMP4 and TGFb, valvular endocardial-like progenitor cells (VEPs) could be efficiently generated from hPSCs under the chemically defined conditions in 9-12 days.

### Comparative transcriptome assay of hPSC-derived VEPs and the primary VECs

Next, a comparative transcriptome analysis was performed to understand to what extent hPSC-derived VEPs resemble the genuine VECs. HPSC-derived VEPs at different time points (day 6 hPSC-derived endocardial cells under the combined treatment with BMP4 and TGFb for 1, 3, 6 days), and the primary VECs freshly isolated from normal human aortic valves of different ages (7-, 9-, 19-, 20-, 30-, 40-, 43-, 50-, 51- and 59-year-old) were used for RNA sequencing (RNA-seq) and the subsequent bioinformatics analysis (Figure 4A) (Table 1). Human ESC H9 cell line (H9), Human foreskin fibroblast cells (HFF), human valvular interstitial cells (hVICs), human umbilical vein endothelial cells (HUVEC) and aortic endothelial cells (HAEC) were also included and used as controls.

**Figure 4.**
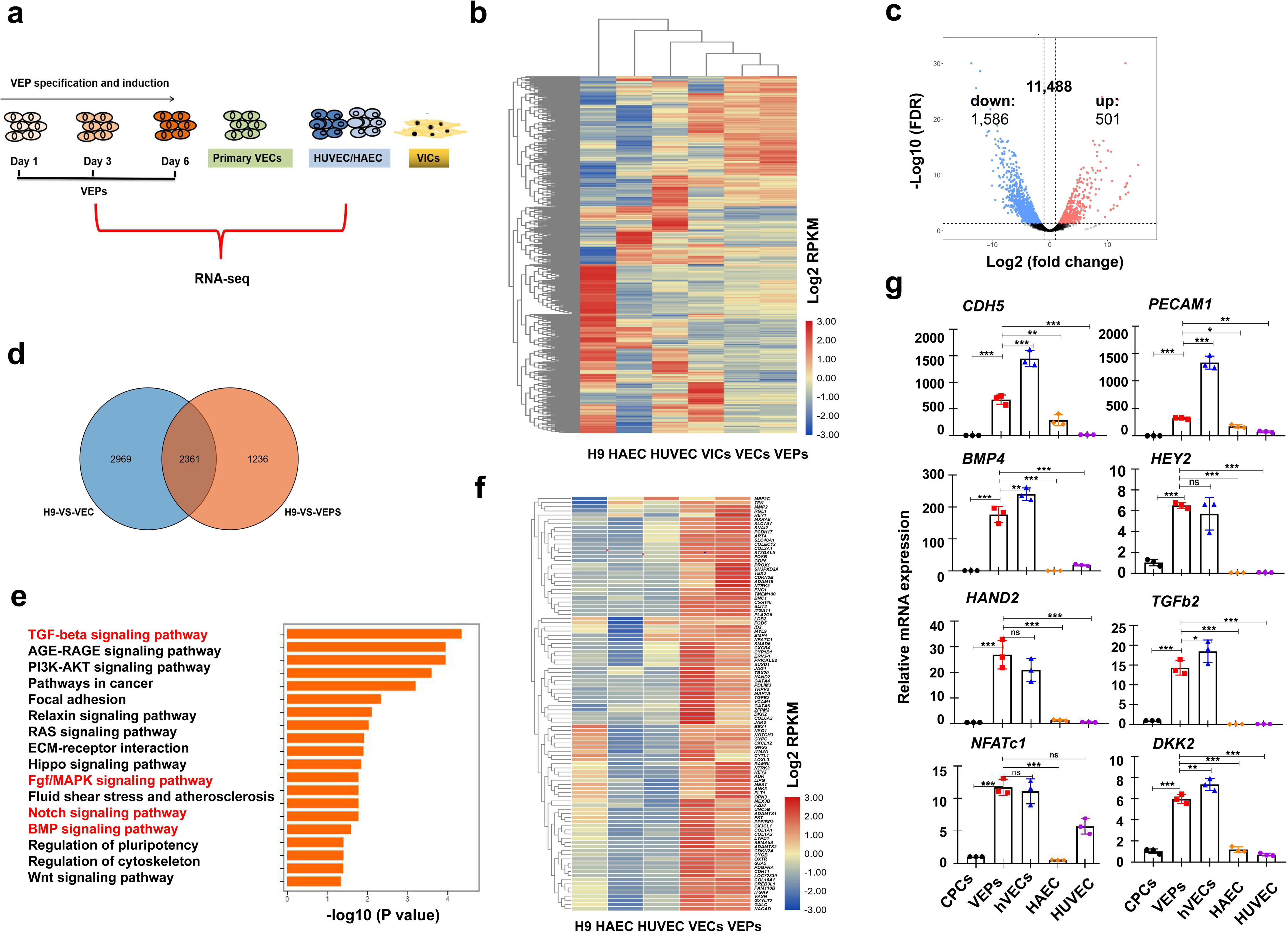
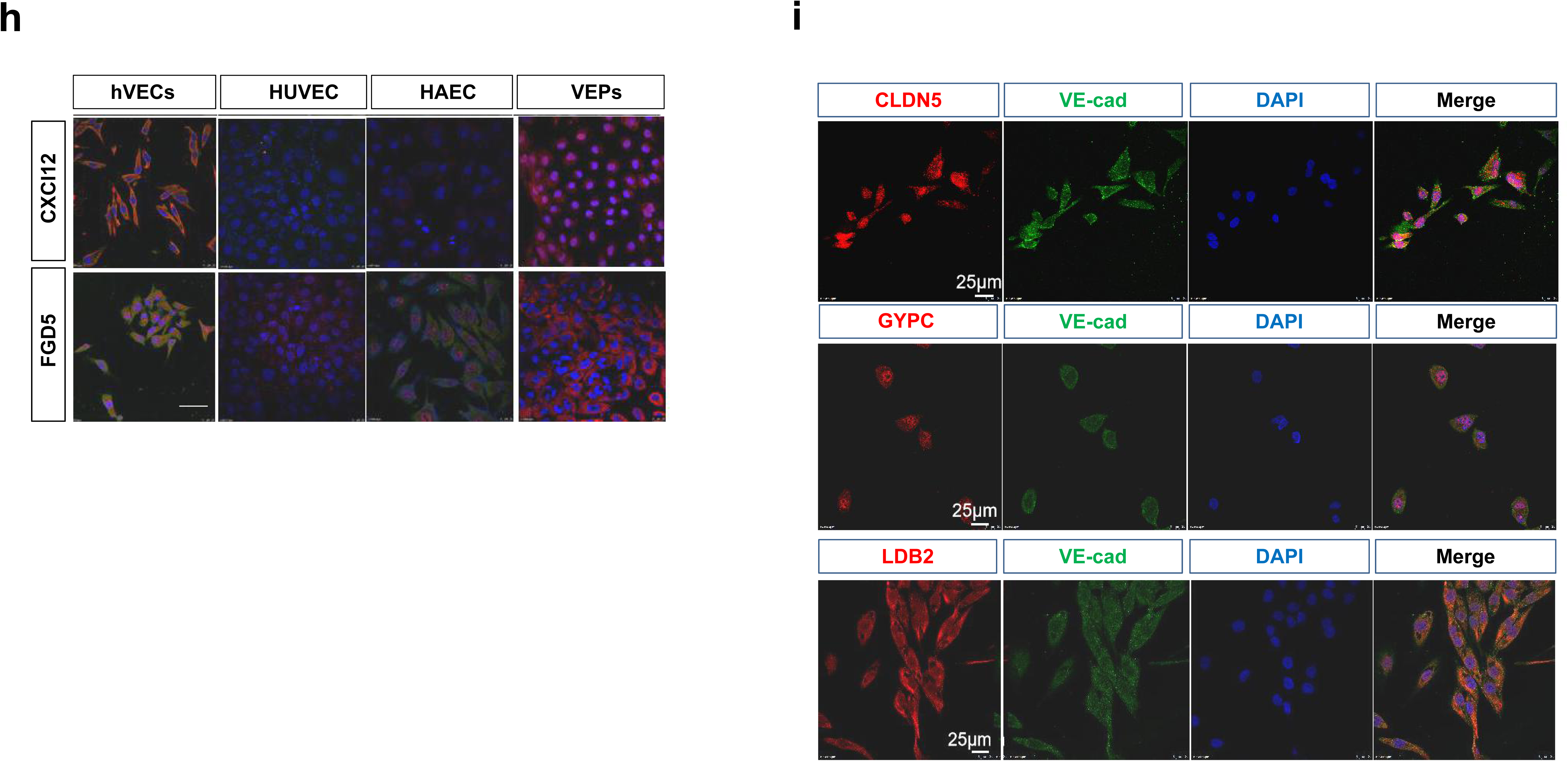
Transcriptome comparison between the VEPs and isolated VECs a. A cartoon showing the design of RNA-seq experiments. hPSC-derived VEPs from three time points (day 1, 3 and 6 post-treatment with VEGFA+ TGFb+ BMP4) were sequenced, and each time points had three experimental repeats. For the primary VECs, the sampling information was shown in Table 1. **b** Heat map showing the similarities among H9, HAEC, HUVEC, the primary VICs from 7-year old, the primary VECs from 7-year old and day 9 hPSC-derived VEPs; Note that day 9 hPSC-derived VEPs are closer to the primary VECs at a global level than H9, HAEC, HUVEC and the primary VICs. Similar results were obtained when using the primary VICs isolated from 10-, 20-, 30-, 40- and 50-year old. **c** Volcano plot showing the similarity between hPSC-derived VEPs and the primary VECs from 7-year old. **d** The pie chart showing the shared genes that are highly expresed in both hPSC-derived VEPs and the isolated primary VECs of 7-year old. **e** KEGG analysis of the shared 218 genes between hPSC-derived VEPs and the primary VECs, showing the enrichment of TGF-β. BMP and Notch signaling pathways. **f** Heat map showing the top 100 genes that are highly expressed in hPSC-derived VEPs and the primary VECs but lowly expressed in H9, HAEC and HUVEC. **g** qPCR validation of the selected VEGs that are highly-expressed in hPSC-derived VEPs and the primary VECs but lowly expressed in hPSC-derived CPCs, HUVEC and HAEC. **h** IF staining showing the expression of FGD5 and CXCL12 in the primary VECs and day 9 hPSC-derived VEPs, but not in HUVEC and HAEC. **i** IF staining showing the expression of GLDN5 GYPC and LDB2 in the primary VECs of 7-year old. Scale bar: 100 μm. All experiments were repeated 3 times. Significant levels are: \**p*< 0.05; \*\**P*< 0.01; \*\*\**P*< 0.001. Shown are representative data.

**Table 1.**
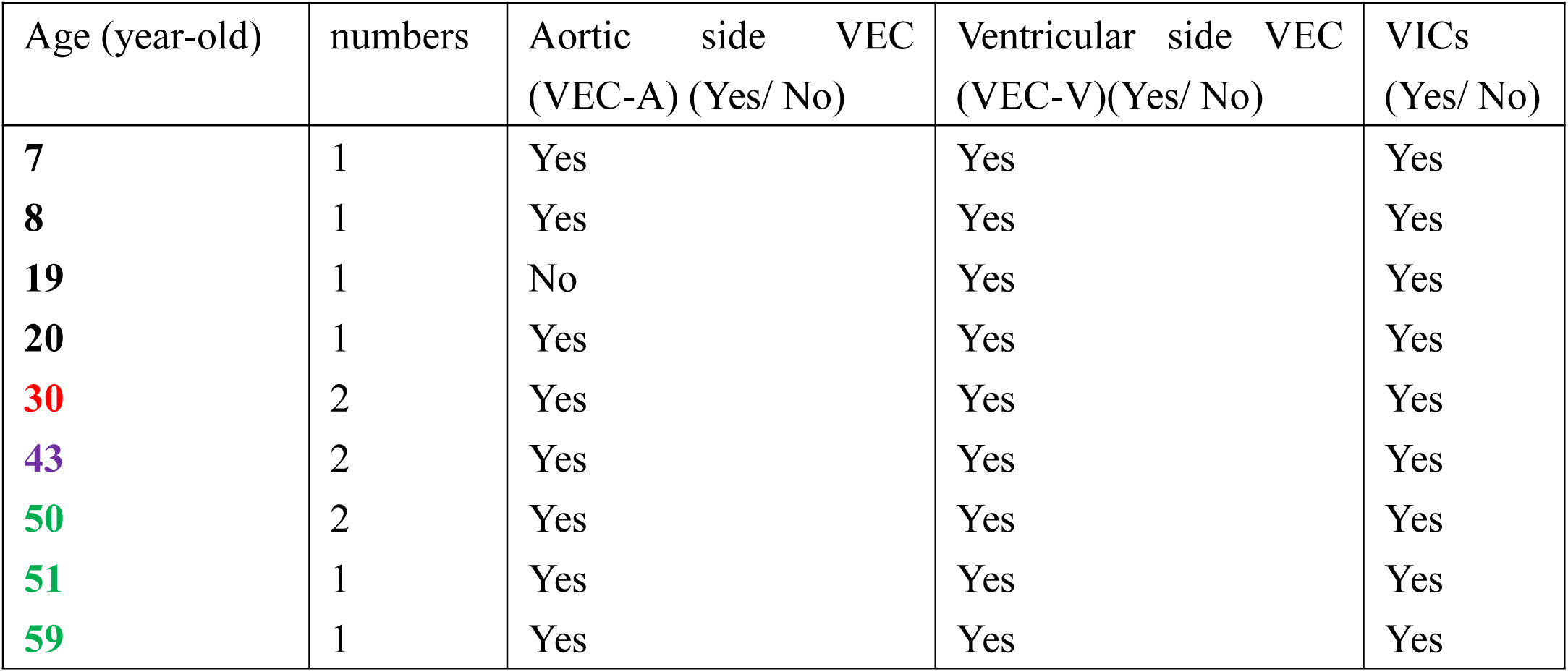
The primary VECs isolated from normal human aortic valves

The RNA-seq results of the primary hVECs of different ages showed that the definitive endothelial markers *CDH5* and *PECAM1* were highly expressed (FPKM> 50) while the fibroblast markers *MMP1/3* and *POSTN* as well as cardiomyocyte markers *TNNT1/2* were barely expressed (FPKM< 2). IF results confirmed that almost all cells were CD144 positive. Thus, we determined that the isolated primary hVECs were of high purity (or rarely contaminated by VICs or other cardiac cell types). An average of 16,000 genes in the primary hVECs were identified, which was similar to that of HUVEC (Supplementary Figure 4A). Of which, approximately 6,000 genes were expressed at higher levels (FPKM> 10); 6,000 genes were expressed at medium levels (FPKM 1-10); and the remainder genes were barely or lowly expressed (FPKM< 1). As shown in Figure 4B, heat map analysis of the normalized RNA-seq data showed that day 9 hPSC-derived VEPs were globally more similar to the primary VECs of different ages than HUVEC, HAEC, hVICs and H9 (Figure 4B; Supplementary Figure 4B). The vast majority of genes (11,488 out of 13,575= 85%) were expressed at similar levels between day 9 hPSC-derived VEPs and the primary hVECs, albeit 2,087 (∼15%) differentially expressed genes (DEGs) were identified (501 genes were up-regulated and 1,586 down-regulated; *P*< 0.05) (Figure 4C). Next, we asked how many highly-expressed genes (FPKM> 10) were shared by day 9 HPSC-derived VEPs and the primary hVECs. We found that a total of 2,361 genes were expressed at similar levels, representing approximately 44% and 65% of total highly-expressed genes of hPSC-derived VEPs and the primary VECs, respectively (Figure 4D). This further supported that HPSC-derived VEPs and the primary hVECs exhibited considerable similarities in gene expression.

To understand the possible genetic signature of hVECs, a comparative transcriptome analysis was performed for the primary hVECs and HUVEC/HAEC to identify VEC-enriched genes (Supplementary Figure 4B). We have identified more than 400 hVEC-enriched genes: highly-expressed (FPKM> 10) in hVECs of 7-year old but lowly-expressed in HUVEC/HAEC (FPKM< 5). Of which, 218 genes (53%) were also highly expressed in day 9 hPSC-derived VEPs, which again supported that hPSC-derived VEPs resembled the primary hVECs. GO (Gene Ontology) analysis of hVEC-enriched genes showed that they were highly related to terms such as cardiovascular and valve development (Supplementary Figure 4C). KEGG pathway enrichment analysis revealed that TGF-β, Notch and FGF/Erk signaling pathways were highly represented (Figure 4E), suggesting that hVECs require robust expression of genes related to these signaling pathways. This implied that enhancing FGF, Notch and TGF-β signaling may facilitate the derivation of valve endocardial-like cells (Hulin et al., 2019; Neri et al., 2019). In this context, the data supported our rationale and method that were utilized for differentiating hPSC into the VEP cell types. Remarkably, KEGG analysis also revealed the enrichment of genes related to fluid shear stress and atherosclerosis, including *ID2*, *BMP4*, *TBX*3, *JAK*2 and *FZD8*. It is increasingly realized that fluid shear stress from blood flow acting on the VECs critically regulates valve sclerosis (Holliday et al., 2011). In future, it will be interesting to investigate whether dysfunction of these genes in hVECs mediates valve sclerosis.

To further understand the genetic signature of hVECs, we screened the top 100 genes that were highly expressed in both the primary hVECs and hPSC-derived VEPs but lowly expressed in HUVEC/HAEC. Heat map analysis was performed and used to display these genes (Figure 4F; Supplementary Figure 4D). This included the known VEGs such as *TGFb2*, *TBX2/3*, *TBX20, GATA4*, *THSD7*, *PROX1*, *HAND2*, *NOTCH1/3* and *HEY1/2,* and new candidate marker genes, such as *BMP4*, *GJA5*, *FGD5*, *EDN1*, *CXCL12*, *CLDN5*, *LDB2*, *CDH11*, *PDGFRa*, *GYPC*, *ISLR*, *HMCN1* and *TSPAN8*. Of the new-identified genes, a few have been implicated in valve endocardial formation. For instance, the chemokine CXCL12 and its receptor CXCR4 are localized to the endocardial cells of the OFT and atrioventricular cushions, and disruption of this signaling causes cardiac defects (Sierro et al., 2007). CDH11 and PDGFRa has been shown to be expressed in valve progenitor cells (El-Rass et al., 2017). Endocardium expressed HAND2 has been shown to control endocardium maturation via regulation VEGF signaling (VanDusen et al., 2015). Of note, compared to HAEC/HUVEC, both hPSC-derived VEPs and the primary VECs expressed higher levels of genes associated with bone formation, including *BMP4*/*BMP6*/*ID2*, *MMPs*, *ITM2A*, *BGN*, *SMAD6*, *MGP*, *PTN*, *HAPLN1*, *FBN1*, *COL1A1*/*COL3A1*, *OSTF1*, *CTSK*, *GALC, ART4, CADM3* and *MYL9*, and genes associated with antioxidative activities, including *vWF*, *GPX3*, *NOS3*, *PRDX2*, *SELP* and *TNFSF4* (Simmons et al., 2005). The qRT-PCR analysis of the selected genes confirmed the RNA-seq results (Figure 4G; Supplementary Figure 4E-G). To validate some of the new candidate markers, IF staining was employed to validate their protein expression. The results showed that CXCL12 and FGD5 were expressed in both the primary hVECs and hPSC-derived VEPs, but barely expressed in HUVEC/HAEC (Figure 4H). Similar results were observed for CLND5, GYPC and LDB2 (Figure 4I).

Both venous (*EPHB4* and *NR2F2*) and arterial markers (*NOTCH1* and *CXCR4*) was abundantly expressed in the primary hVECs and hPSC-derived VEPs (Figure 4F), suggesting that hVECs might be not biased to any of the two vascular endothelial types. Fibroblast marker genes such as *COL1A1, MMP2* and *BGN* were found to be expressed in both hVECs and hPSC-derived VEPs (Supplementary Figure 3B; 4F). It has been reported that the primary VECs isolated from human aortic valves co-express both endothelial and fibroblast markers (Holliday et al., 2011). Thus, this might not be due to the EndoMT of hPSC-derived VEPs.

One unique feature of hVECs is that they are constantly exposed to the oscillatory shear flow. We therefore examined the mRNA expression of oscillatory shear-related genes such as *EDN1*, *BMP4*, *CTSK*, *PPARG* and *THBS1* (Holliday et al., 2011). The RNA-seq results showed that these genes were expressed at significantly higher levels in the primary VECs than HUVEC/HAEC. Next, the expression levels of oscillatory shear-related markers were compared between the primary hVECs and hPSC-derived VEPs. We found that they were expressed at significantly lower levels in hPSC-derived VEPs than the primary VECs. Similar results were observed for general shear stress response genes such as *KLF2*, *CAV1*, *NOS3* and *ICAM1*. The qRT-PCR analysis of selected markers confirmed the RNA-seq results (Supplementary Figure 3B, 4F-H). These observations suggest that hPSC-derived VEPs might be in an immature state, presumably due to lack of the hemodynamic stimulus that are exposed to human valve endothelial cells in normal physiological conditions (see discussion below).

### BMP4 and TGFb promote VEP fate by enhancing *NFATc1* and *ETV2*

So far, we have shown that the crosstalk among BMP, FGF and TGF-β signaling pathways is important for hPSC differentiation into the VEPs. To further understand the underlying mechanisms, hPSCs were differentiated into the VEPs in the presence and absence of BMP signaling inhibitor LDN and TGF-β signaling inhibitor SB431542 (SB). The qRT-PCR results have shown that the mRNA expression of VEGs such as *JAG2*, *HEY2*, *TBX2* and *NFATc1* were significantly reduced in the presence of LDN and SB (*P*< 0.05; Supplementary Figure 3A, 3B; Figure 3B). Consistently, the flow cytometry results showed that the percentage of NOTCH1 positive cells was greatly reduced in day 9 cell cultures treated with LDN and SB compared to the control (30% in treated vs 80% in control) (Figure 5A). Notch-related genes *JAG1, HEY1, TBX2/3* and *SNAI1,* are well-known direct targets of TGF-β/BMP signaling (Garside et al., 2013; Hoogaas et al., 2007; Morikawa et al., 2011; Zawadil et al., 2004). Thus, the combined treatment with BMP4 and TGFb may induce the VEP fate by activating these Notch-related factors, which was in line with a recent report (Papoutsi et al., 2018).

**Figure 5.**
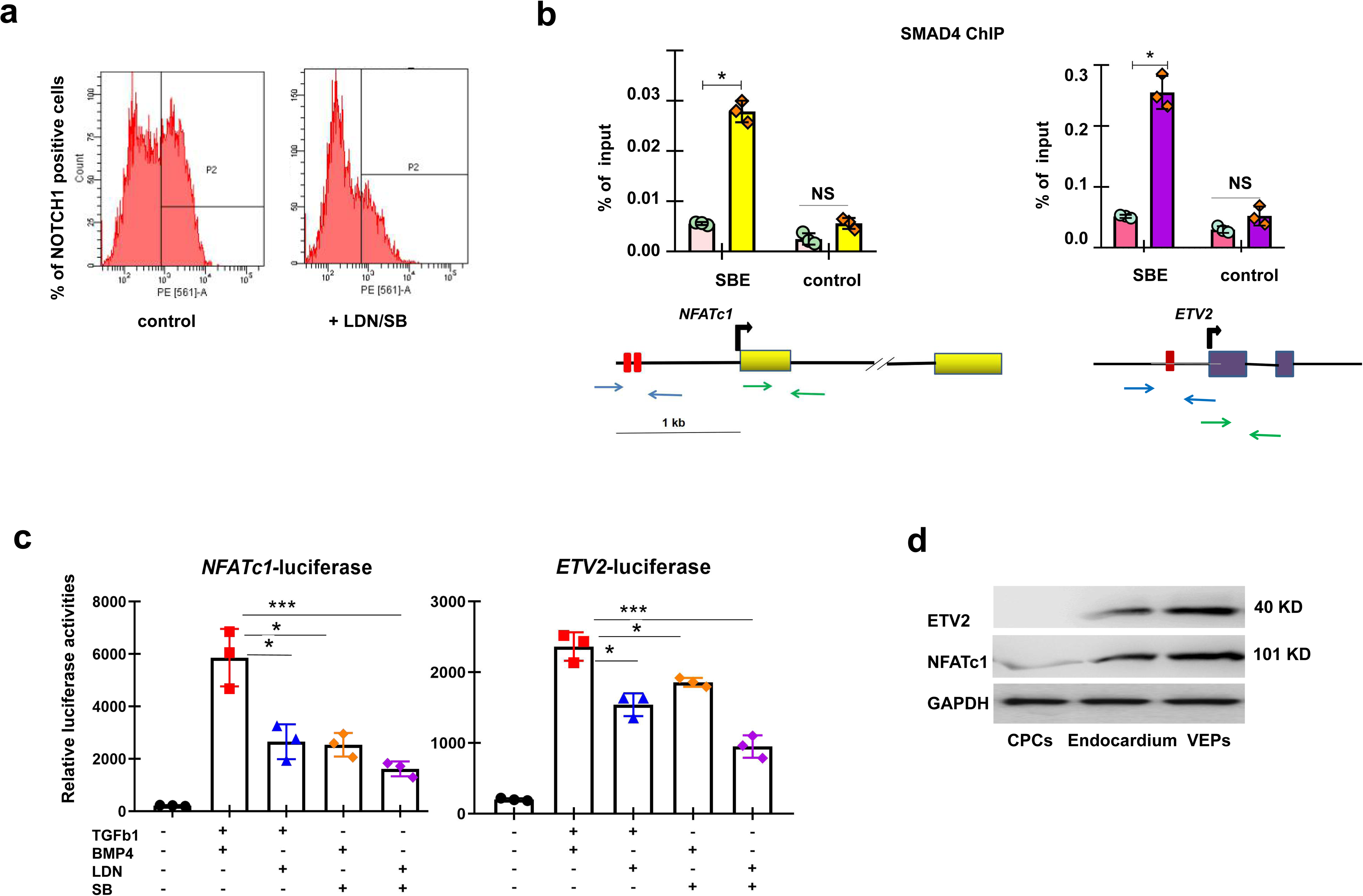
BMP and TGF induce VEP fate by promoting NFATc1 and ETV2 a. Flow cytometry analysis of day 9 hPSC-derived VEPs in the presence and absence of SB and LDN, showing the percentage of NOTCH1 positive cells. **b** ChIP experiments showing the enrichment of SMAD4 at *NFATc1* (left) and *ETV2* (right) promoter regions. **c** Luciferase reporter assay showing *NFATc1* and *ETV2* promoter activities in the presence and absence of LDN and SB. **d** WB analysis of the indicated antibodies for hPSC-derived CPCs, endocardial cells and VEPs. All experiments were repeated 3 times. Significant levels are: \**p*< 0.05; \*\**P*< 0.01; \*\*\**P*< 0.001. Shown are representative data.

A key feature of valve endocardial cells is the sustained expression of NFATc1 and ETV2 (Wu et al., 2012). NFATc1 is the key regulator for the induction, maintenance and fate determination of valve endocardial cells (Wu et al., 2012). It has been shown that *NFATc1* expression levels remain high in the pro-valve endocardial cells, which is important for the maintenance of VEP identity in mice (Wu et al., 2011). Mouse *Nfatc1* gene contains SMAD binding elements (SBEs) in enhancer regions that are important for its pro-valve endocardial cell specific expression (Zhou et al., 2004). SMAD transcription factors, including the R-SMADs and the common SMAD (SMAD4), are the mediators of signal transduction by TGF-β superfamily members which include BMP and TGF-β (Mandal et al., 2016). Etv2, the earliest restricted regulator of endothelial development which is thought to function upstream of *NFATc1*, plays key role in initiating endocardial differentiation in different animal models (Nakano et al., 2016; Palencia-Desai et al., 2011; Puceat, 2013). In fact, *ETV2* and *NFATc1* were among the earliest markers that were induced by the combined treatment with BMP4 and TGFb (Figure 3C). We speculated that the combined treatment with BMP4 and TGFb may promote VEP specification through modulating the expression of *NFATc1* and *ETV2*. Our hPSC VEP differentiation system provides us with a unique system to test this hypothesis in vitro. First, we identified two SBEs composed of the sequence CAGACA in the proximal promoter of the human *NFATc1* gene, and one SBE in the proximal promoter of the human *ETV2* gene (Figure 5B). ChIP-PCR analysis was performed to validate SMAD4 binding at the gene promoters. The ChIP-PCR results showed that SMAD4 was indeed enriched at both *NFATC1* and *ETV2* promoter regions containing the SBEs in day 9 hPSC-derived VEPs (Figure 5B). Next, luciferase reporters containing the -2kb *NFATc1* and *ETV2* promoters were constructed and subsequently transfected into HEK293T cells in the presence and absence of BMP4 and TGFβ1 or the inhibitors of the respective signaling pathways. The results of the luciferase reporter analysis showed that treatment with BMP4 and TGFβ1 strongly induced while treatment with LDN and SB repressed the luciferase activities (Figure 5C). Finally, WB analysis was performed to detect the protein levels of ETV2 and NFATc1 during hPSC differentiation into the VEPs. The results showed that the expression of two transcription factors was gradually increased (Figure 5D). When LDN and SB were applied, the expression levels of ETV2 and NFATc1 were significantly reduced (Supplementary Figure 5A).

Another interesting observation is the activation of markers related to bone formation, particular the BMP related genes. *BMP2/4/6* and BMP target genes *TBX2/3* were expressed at significantly higher levels in hVECs and hPSC-derived VEPs than HAEC/HUVEC (Figure 3C; 4F). Two SBEs composed of the sequence CAGACA/TGTCTG were identified in the proximal promoters of human *BMP2/4/6* genes. The ChIP-PCR analysis confirmed that SMAD4 was enriched at *BMP2/4/6* promoter-proximal regions containing the SBEs in day 9 hPSC-derived VEPs (data not shown), which strongly suggests that the combined treatment of BMP4 and TGFb promotes the expression of *BMP2/4/6*.

As shown in Figure 3C, the dynamic expression patterns of the Notch related genes such as *JAG1*/*HEY2*/*NOTCH3/DLL4* were well correlated to other VEGs such as *NFATc1*, *GATA4*, *TEK* and *MEF2*C. This strongly suggested that Notch activation might be a part of VEP specification. However, it has been established that TGF/Notch activation is incompatible to long term EC maintenance as it promotes the EMT process (Wang et al., 2013; Zawadil et al., 2004). In line with this, we found that the EndoMT-related gene *SNAI1* and the mesenchymal genes *COL1A1* and *COL3A1* were significantly induced day 3-4 post-treatment (Supplementary Figure 3B). This suggested that hPSC-derived VEPs were prone to undergo EndoMT to become mesenchymal-like cells when cells were exposed to prolonged treatment with BMP4 and TGFb. It has been shown that functional inactivation of JAG1 or Notch inhibits the EMT induced by TGF/SMAD signaling (Zawadil et al., 2004). We speculated that Notch inhibition by DAPT may stabilize hPSC-derived VEPs by suppressing the EndoMT. To test this hypothesis, day 9 hPSC-derived VEPs were cultured in EBM2 medium in the presence or absence of DAPT. In the absence of DAPT, hPSC-derived VEPs could be maintained up to 2 passages before eventually adopted a fibroblast-like morphology, presumably due to the EndoMT (Supplementary Figure 5B). By contrast, the VEPs could be passaged up to 4-5 times in the presence of DAPT while remaining with the cobblestone EC morphology. The qRT-PCR results showed that passage 3-4 hPSC-derived VEPs still abundantly expressed *PECAM1, CDH5* and *NFATc1*, although the expression levels were reduced to some extent compared to the controls (Supplementary Figure 5C). We also investigated the effect of inhibition of TGF-β signaling by the addition of SB. Unfortunately, SB administration quickly led to fibroblast-like morphology of the VEPs (Supplementary Figure 5D). As extracellular signal-regulated kinase (ERK) has also been implicated as mediator of TGF-β-induced EMT, and ERK inhibition can block it (Lamouille et al., 2014). We asked whether treatment with ERK inhibitors U0126 or PD could be beneficial to maintain the VEP phenotype in the culture plates. We seeded day 9 hPSC-derived VEPs at a density of 1× 10^4^ cells/6-well, and cultured them with EBM2 in the absence or presence of the ERK inhibitors. In the absence of PD, hPSC-derived VEPs were proliferative and maintained the EC morphology. In the presence of PD, however, the cells failed to proliferate although the morphology was well-maintained (Supplementary Figure 5D).

Taken together, we tentatively concluded that the combined treatment with BMP4 and TGFb may induce the VEP fate by enhancing a cohort of genes, including the early endocardial marker genes *NFATc1* and *ETV2,* Notch-related genes *JAG1* and *HEY2*, as well as BMP-related genes *BMP2/4/6*. Notch activation is likely a part of VEP induction. After the specification of VEPs, functional integration among TGF-β, BMP and Notch signaling leads to activation of *JAG1/HEY2/SNAI1* which promotes EndoMT of the transient VEP cell types in vitro.

### Functional characterization of hPSC-derived VEPs

Our final protocol for hPSC differentiation into VEPs is illustrated in Figure 6A. On average, 3 million CPCs could be obtained from 1 million hPSCs in 3 days. After additional 6-9 days of differentiation via an intermediate endocardium stage, 10-20 million VEC-like progenitor cells can be generated depending our hPSC lines. HPSC-derived VEPs could be passaged up to 4 times with commercial EBM2 medium before eventually adopting a fibroblastic morphology (Supplementary Figure 6A).

**Figure 6.**
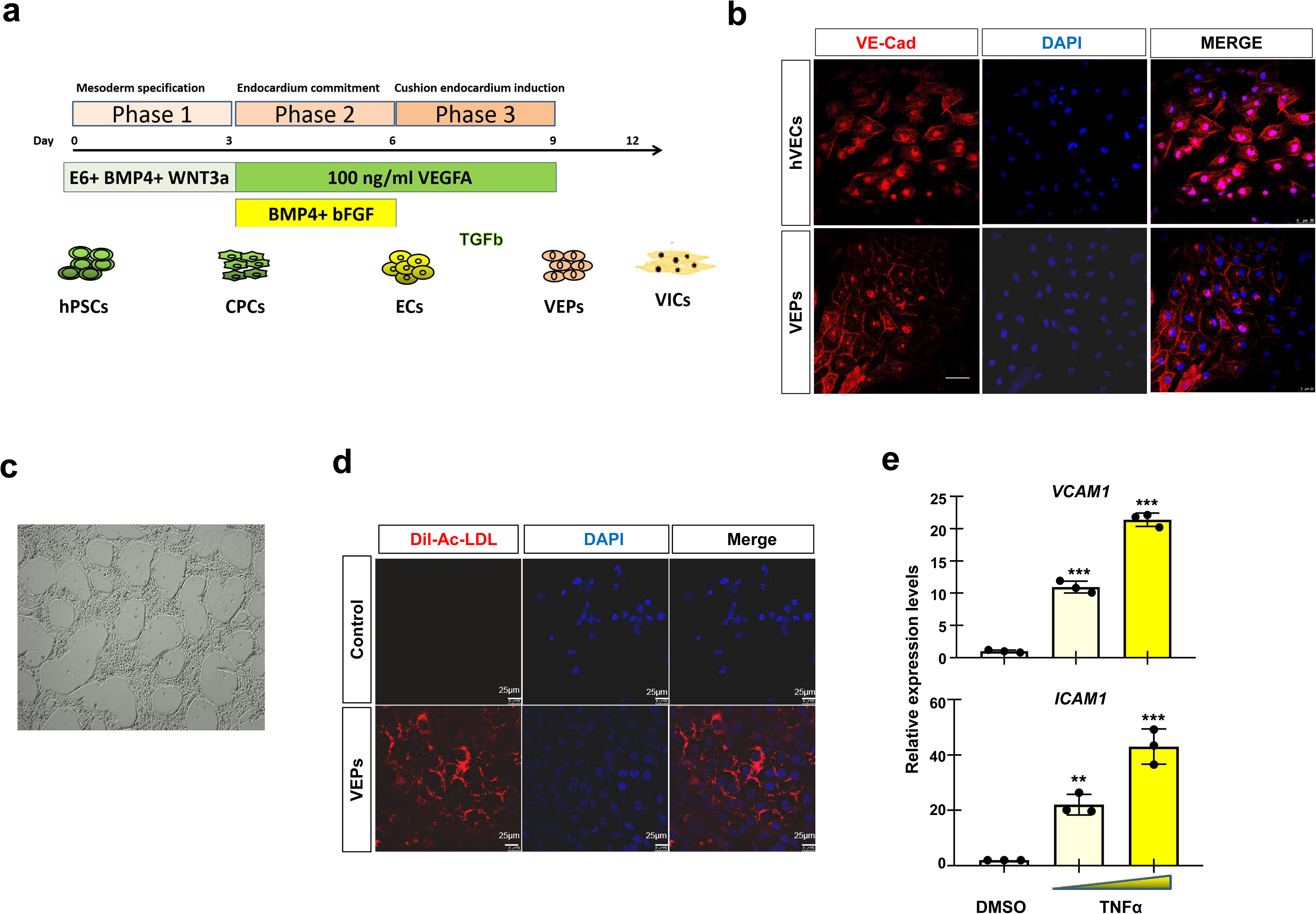
Functional characterization of VEPs a. Cartoon depicting hPSC differentiation to VEPs for 9-12 days. **b** IF staining showing that hPSC-derived VEPs express VE-cad similar to the primary VECs. Scale bar: 25 μm. **c** Tube formation assay of day 9 hPSC-derived VEPs. **d** Fluorescence signals showing that hPSC-derived VEPs could uptake Ac-LDL. **e** The qPCR analysis of *VCAM1* and *ICAM1* genes for hPSC-derived VEPs in the presence and absence of TNFa. All experiments were repeated 3 times. Significant levels are: \**p*< 0.05; \*\**P*< 0.01; \*\*\**P*< 0.001. Shown are representative data.

In addition to cell type-specific marker characterization, functional assays are important determinants of cell identity and cellular maturation (Glaser et al., 2011). We therefore investigated whether hPSC-derived VEPs functionally resembled the primary VECs. A hallmark of ECs is the uptake of LDL both for normal metabolism and under pathologic conditions (Holliday et al., 2011). HPSC-derived VEPs were split at a 1:3 ratio and re-plated on Matrigel, and fluorescent-acetylated LDL (Dil-Ac-LDL) uptake assay was performed. HPSC-derived VEPs adopted an EC cell morphology and formed vascular network-like structures on Matrigel, and abundantly expressed the endothelial markers CD31 and TIE2 as well as the newly-identified candidate markers CXCL12 and FGD5, similar to the primary VECs (Figure 6B, C; Supplementary Figure 6B, C). To assess the LDL uptake, day 9 hPSC-derived VEPs and the primary VECs were incubated with Alexa Fluor 594 conjugated to acetylated LDL (Ac-LDL) for 2 hours. HPSC-derived VEPs were able to robustly incorporate Dil-Ac-LDL and exhibited relatively homogeneous uptake of LDL as indicated by the red fluorescence in each cell (Figure 6D).

It is increasingly realized that hVECs may play a critical role in the pathogenesis of valvular heart disease, and regulate vascular tone, inflammation, thrombosis, and vascular remodeling (Hinton and Yutzey, 2011). Calcific aortic valve sclerosis involves inflammatory processes and occurs on endothelialized valve leaflets (Simmons et al., 2005; Holliday et al., 2011). We investigated whether hPSC-derived VEPs could response to pro-inflammation stimulus. To this end, hPSC-derived VEPs were seeded in 6-well plates at a density of 4,000/cm^2^ and grown to confluence using EBM2 media. Confluent monolayers were incubated with 0.1 or 10 ng/ml recombinant TNFα for 4 hours. We found that *VCAM1* and *ICAM1* were significantly induced by TNFα, in a dose dependent fashion (Figure 6E). These data suggested that hPSC-derived VEPs could increase the expression levels of cell surface markers *VCAM1* and *ICAM1* in response to inflammatory stimulus.

In summary, our hPSC-derived VEPs exhibited similar morphological, molecular and functional features to that of the primary VECs.

### VEPs become VIC-like cells through EMT

During valvulogenesis, valvular interstitial cells (VICs) are mainly derived from endocardial cushion cells through an EndoMT process that is regulated by several signaling pathways, including TGF-β, BMP and FGF signaling pathways (Combs and Yutzey, 2009; Hinton and Yutzey, 2011; Lincoln et al., 2006). We therefore investigated whether hPSC-derived VEPs could be converted to VIC-like cells through inducing EndoMT, by treating the VEPs with higher concentration of TGFb and bFGF over a time course of 6 days. The treated cells were collected every 1.5 days and analyzed as described below (Figure 7A; Supplementary Figure 7A). We found that hPSC-derived VEPs quickly lose the cobble-stone morphology. In stead, the fibroblastic morphology was formed (Figure 7B). The qRT-PCR results showed that after 3 days treatment with bFGF and TGFb, VIC-type markers such as *POSTN ACTA2* and *S100A4*/*FSP1* were strongly induced (Figure 7A). In addition, the expression of EndoMT-related genes such as *SNAI1* and *TWIST1*, BMP target genes *SOX9* and *AGGRECAN* as well as FGF target genes *TENASCIN* and *SCLERAXIS*, was significantly increased (Lincoln et al., 2006) (Supplementary Figure 7B). IF staining and WB analyses of the treated cells confirmed that they abundantly expressed FSP1, MRTF-A and VIM, albeit to a lesser levels compared to the isolated primary VICs (Figure 7B-C; Supplementary Figure 7C) (Latif et al, 2017). Thus, VIC-like cells were formed when hPSC-derived VEPs were treated with higher concentration of TGFb and bFGF.

**Figure 7.**
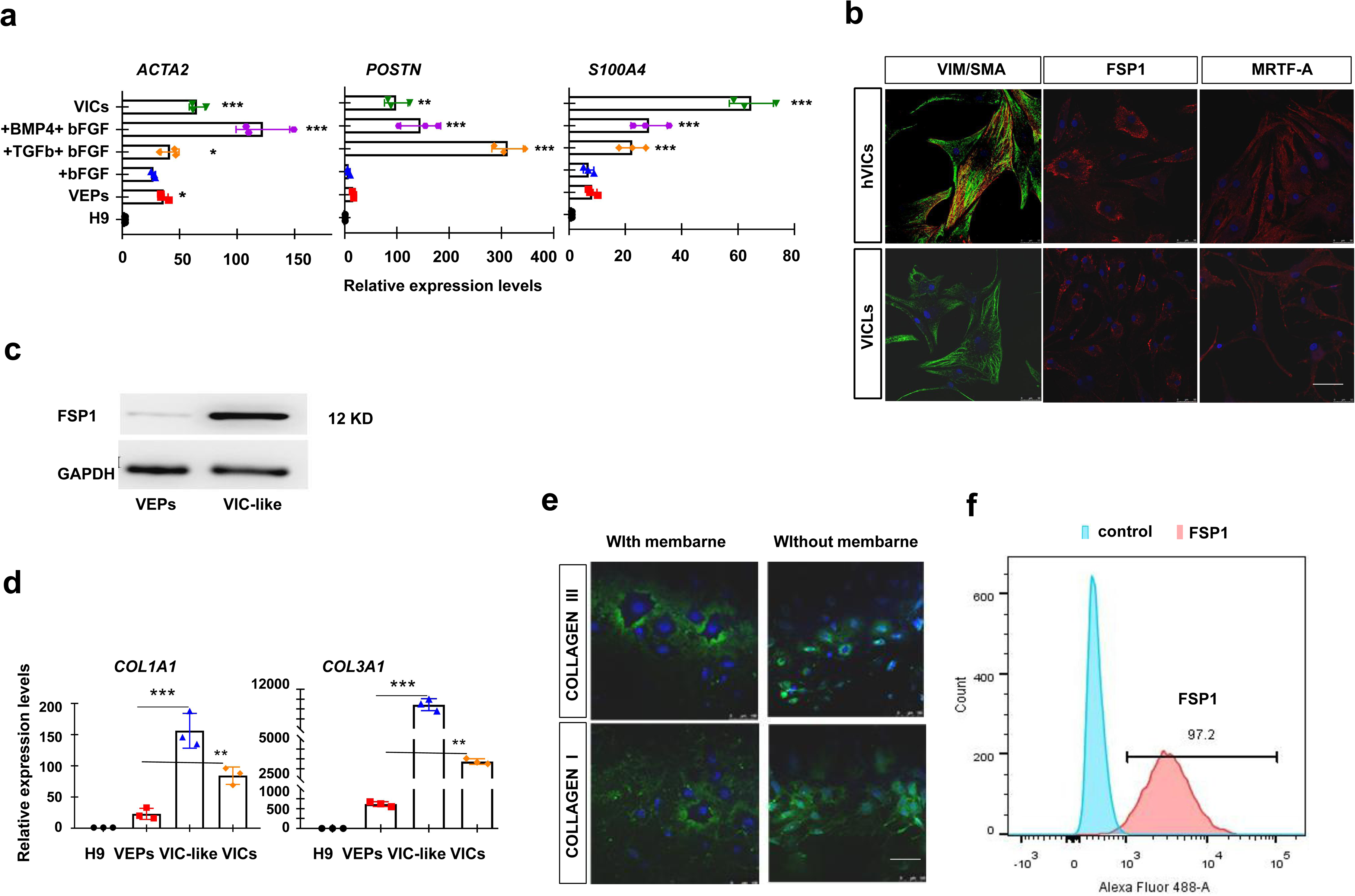
hPSC-derived VEPs to VIC-like cells by inducing EndoMT a. The qPCR analysis of the indicated VIC markers for hPSC-derived VEPs that were treated with the indicated molecules for 3-6 days. **b** IF staining showing that hPSC-derived VIC-like cells expressed the indicated VIC markers, similar to the primary VICs. Scale bar: 100 μm. **c** WB analysis showing the expression levels of FSP1 in hPSC-derived VIC-like cells and VEPs. **d** The qPCR analysis of type I collagen marker *COL1A1* and type III collagen marker *COL3A1 for* hPSC-derived VIC-like cells. **e** IF staining showing the expression of COLLAGEN I and III in hPSC-derived VIC-like cells with and without membrane breaking treatment. **f** Flow cytometry assay showing the percentage of FSP1 positive cells in hPSC-derived VIC-like cells. All experiments were repeated 3 times. Significant levels are: \**p*< 0.05; \*\**P*< 0.01; \*\*\**P*< 0.001. Shown are representative data.

Next, we investigated whether the VIC-like cells expressed extracellular matrix components (ECM). We found that the expression of ECM-related genes such as *COLLAGEN I* (*COL1A1*) and *III* (*COL3A1*) was significantly increased in hPSC-derived VEPs that treated with bFGF and TGFb for 6 days (Figure 7D). To investigate whether VIC-like cells synthesize the ECM, IF staining was performed using antibodies against COLLAGEN I and III. The results showed that the VIC-like cells abundantly expressed COLLAGEN I and III (Figure 7E). Finally, flow cytometry analysis of hPSC-derived VIC-like cells revealed that the vast majority of the cells (> 97%) were FSP1 positive (Figure 7F). Taken together, we concluded that valvular interstitial-like cells can be derived from hPSC-derived VEPs by inducing EndoMT.

### HPSC-derived VEPs seeded on de-cellularized porcine heart valves

De-cellularized bioprosthetic scaffolds, which have been widely used to replace the aortic valve, have exhibited promising results for endogenous endothelialization and mechanical durability (Nejad et al., 2016). We have previously made poly(ethylene glycol) tetraacrylate (PEG-TA) cross-linked de-cellularized porcine aortic valves (PEG-DCVs) (Hu et al., 2013). Although the acellular PEG-DCVs exhibit improved mechanical and anti-calcification properties when compared to the glutaraldehyde crosslinked counterparts, the poor clinical long-term result remains unsatisfactory likely due to the early tissue degeneration (Zhou et al., 2013). Thus, the next generation tissue engineered heart valves (TEVs) that overcome the shortcomings of the current mechanical and bioprosthetic valves are highly demanded.

To initially evaluate whether hPSC-derived VEPs could be used as the potential seed cells for the next generation TEVs, we studied the growth and adherent properties of the VEPs seeded on the de-cellularized porcine aortic valves (DCVs). We seeded a total of 4×10^5^ hPSC-derived VEPs on the DCVs in the presence of 100 ng/ml VEGF and 20 ng/ml bFGF, and the cells were monitored for a time course of 7 days. We found that 1 day after seeding, about 50% of hPSC derived VEPs attached to the surface, similar to HUVEC as revealed by IF staining of F-actin antibodies (Figure 8A). The attached cells expressed the general EC markers CD31 and VE-cad (Supplementary Figure 8A, B). Cell proliferation assay by EdU staining showed that hPSC-derived VEPs were highly proliferative, with 40% of cells being EdU positive at day 3 and 80% at day 4 (Figure 8B). In contrast, apoptosis assay by TUNEL staining showed that TUNEL positive cells were reduced from approximately 19% at day 2 to 4% at day 4 (Figure 8C). Analysis of dynamic growth for a time course of 7 days demonstrated a robust growth of the seeded VEPs on the DCVs (Figure 8D). IF staining of day 7 cells showed that they co-expressed TIE2 and VIM as well as VE-Cad and BMP4 (Figure 8E). BMP4 has been shown to be expressed in hVECs in physiological and shear stress conditions (Holliday et al., 2011).

**Figure 8.**
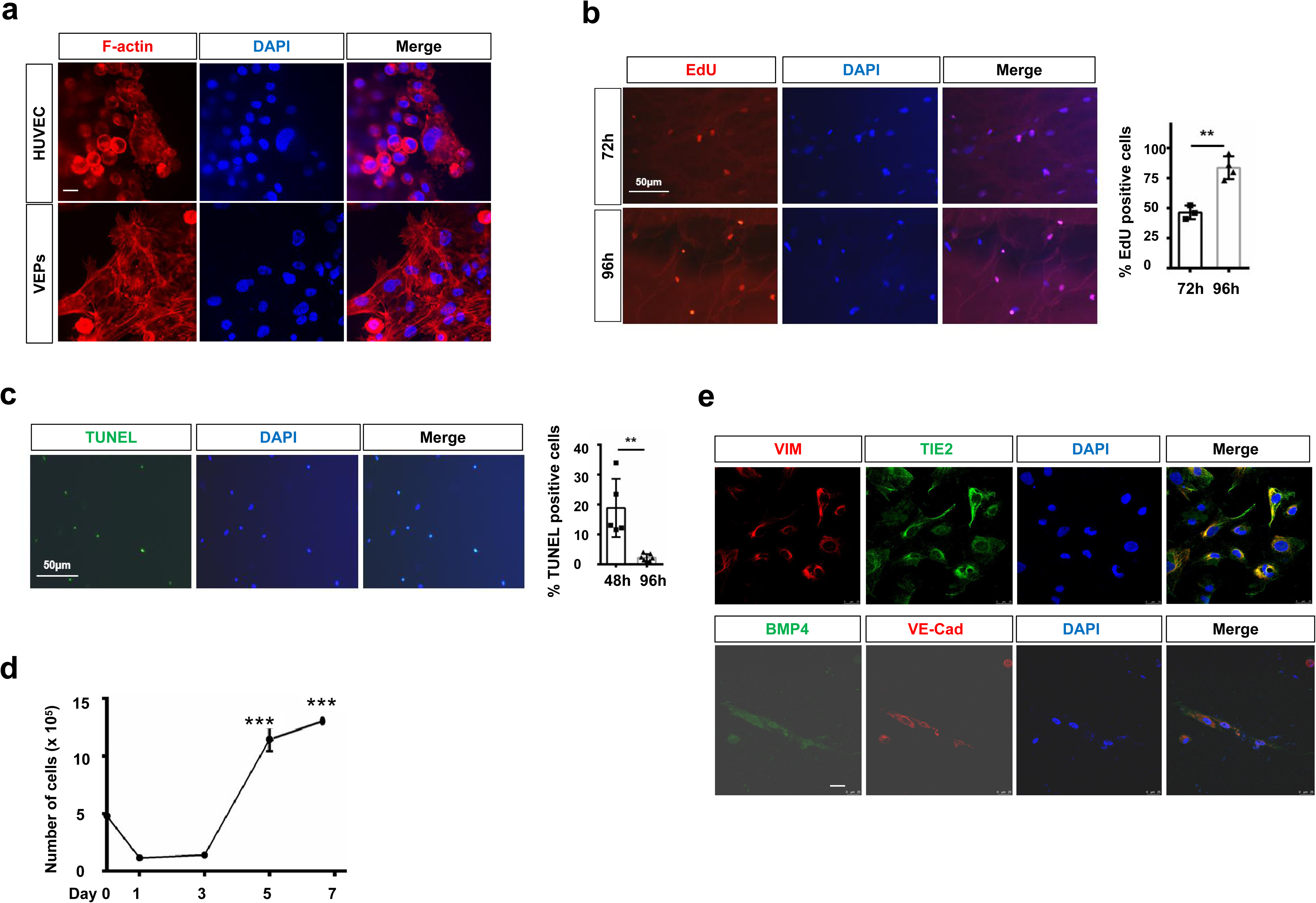
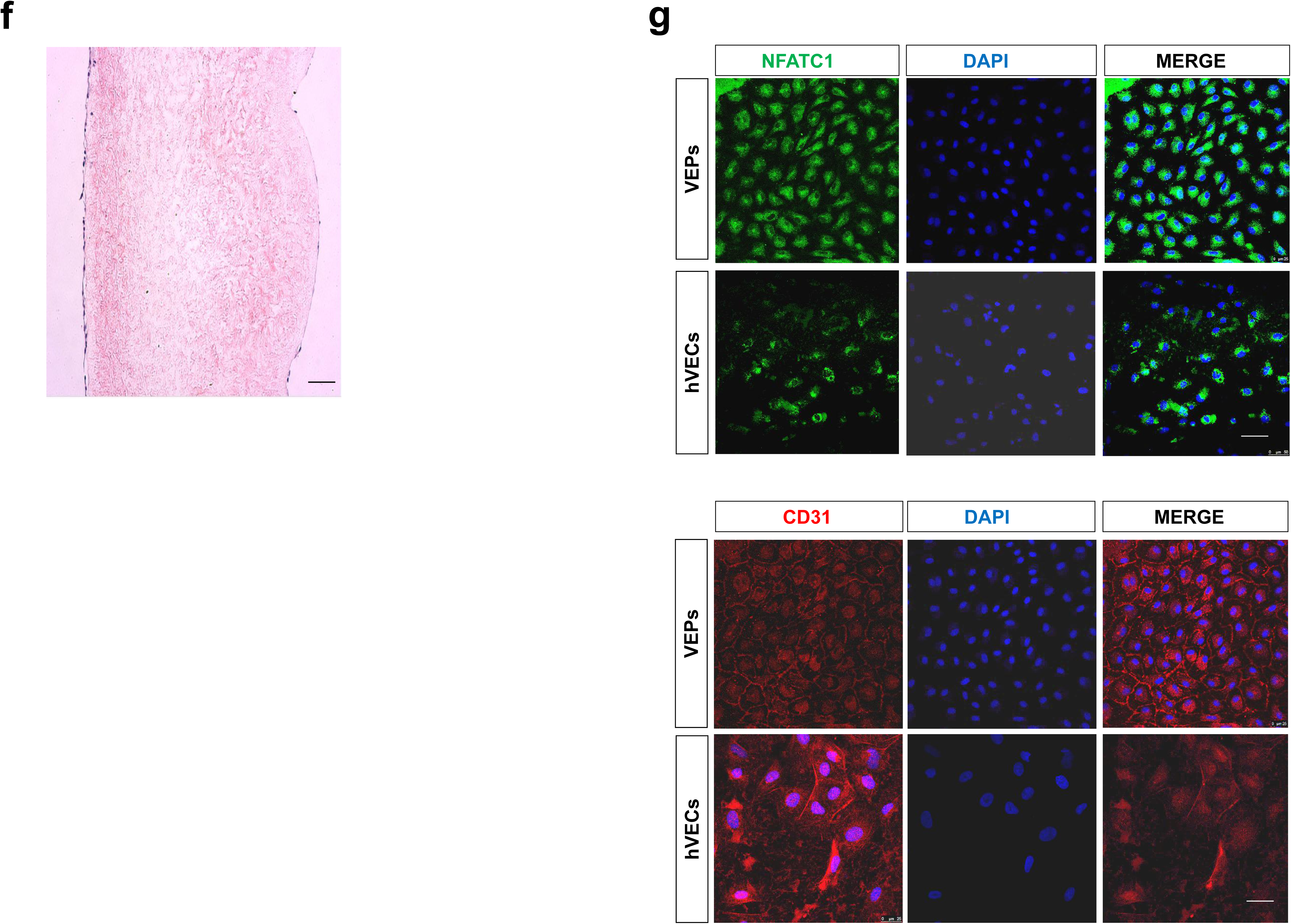
HPSC-derived VECs seeded on de-cellularized porcine heart valves a. IF staining of F-actin showing the morphology of hPSC-derived VECs and HUVEC seeded onto the de-cellularized porcine heart valves. Scale bar: 25 μm. **b** EdU staining of hPSC-derived VEPs seeded onto the de-cellularized porcine heart valves after 72 hours and 96 hours. Right panel: the percentage of EdU positive cells at 72h and 96h. **c** TUNEL assay for hPSC-derived VEPs seeded onto the de-cellularized porcine heart valves. Right panel: the percentage of TUNEL positive cells at 48h and 72h. **d** 7-day growth curve of hPSC-derived VEPs seeded onto the de-cellularized porcine heart valves. **e** IF staining showing CD31, VE-Cad, and VIM signals in hPSC-derived VEPs seeded onto the surface of the de-cellularized porcine heart valves. Bar: 50 μm. **f** The IHC result showing the lining of the DCVs with hPSC-derived VEPs after 3 weeks of co-culturing. Lateral view. Left: aortic side; Right: ventricular side. **g** IF staining showing NFATc1 and CD31 signals in hPSC-derived VEPs that have been seeded on the surface of the de-cellularized porcine heart valves for 3 weeks. Scale bar: 100 μm. All experiments were repeated 3 times. Significant levels are: \**p*< 0.05; \*\**P*< 0.01; \*\*\**P*< 0.001. Shown are representative data.

To understand the long term interaction between hPSC-derived VEPs and the DCVs, we seeded hPSC-derived VEPs at a low density and co-cultured them for 3-4 weeks. The DCVs were washed with pulsatile flow of culture medium before examination. A monolayer of endothelium was formed on the surface of the valves, as revealed by immunohistochemistry (IHC) staining for CD31 (Figure 8F). IF staining of the co-cultured DCVs showed that NFATc1 and CD31 were abundantly expressed in the surface of the DCVs, confirming the IHC result (Figure 8G). Interestingly, IF staining showed that hPSC-derived VEPs appeared more evenly distributed and more proliferative than the primary VECs. These results demonstrated that valve endothelialization occurred after long term co-culture with hPSC-derived VEPs, and that hPSC-derived VEPs exhibited superior proliferative and clonogenic potential than the primary VECs.

Taken together, we concluded that hPSC-derived VEPs could attach, survive and proliferate when seeded on the de-cellularized porcine aortic valves, suggesting that they could have potential implications for modeling valve diseases or use for seed cells for the next generation tissue engineered valves.

## Discussion

In this work, we reported that using a chemically defined xeno-free method, the VEC-like progenitor cells resembling human valve endothelial cells can be rapidly and efficiently generated from hPSCs without a sorting step. HPSC-derived VEPs resemble the primary VECs morphologicaly, molecularly and functionally, and could undergo EndoMT to form VIC-like cells. Furthermore, hPSC-derived VEPs seeded onto the surface of the DCV scaffolds recapitulate key features similar to that of the primary VECs isolated from human cardiac valves. Importantly, endothelialization of the DCVs was observed when hPSC-derived VEPs and the DCVs were co-cultured for 3-4 weeks, which demonstrated that hPSC-derived VEPs could be used as potential seed cells for the next generation TEVs in the future.

A recent study has reported that pre-valvular endocardial cells (HPVCs) can be derived from hPSCs (Neri et al., 2019). However, the method described by Neri and colleagues lacks of sufficient details and the major conclusion was drawn primarily based on comparative transcriptome analysis between hPSC-derived HPVCs and mouse atrio-ventricular canal (AVC) endocardial cells which obviously should be the human AVC endocardial cells or valve endocardial cells. Thus, the developmental biology underlying their hPSC differentiation approach was unclear. Furthermore, the differentiation approach requires cell sorting, the use of mouse fibroblast cells and the knockout serum, which makes the differentiation variable and invokes potential safety problem.

In this work, we describe a chemically defined xeno-free method for generating VEC-like progenitor cells from hPSCs, by a three step-wise differentiation strategy that mimic the embryonic valvulogenesis. First, our finding revealed a key role of FGF signaling in directing hPSC-derived CPCs to an endocardial fate. Studies in mouse, chick and zebrafish have implicated FGF signaling in valvular cell specification and cushion morphogenesis (Marques et al., 2008; Park et al., 2008), and inhibition of FGF signaling resulted in significantly reduced expression of the endocardial markers and defective valvulogenesis (Alsan et al., 2002; Barron et al., 2000). Consistently, FGFs are required for the derivation of cardiac and endothelial cells from hPSCs (Palpant et al., 2017; White et al., 2013; Neri et al., 2019). We extended the previous findings showing that BMP4 administration is beneficial for the derivation of endocardium in the presence of bFGF. We propose that a crosstalk of FGF and BMP signaling may be required for optimal specification of endocardial progenitors, which is in line with a previous study showing that FGF and BMP are both required to induce early cardiac genes in chicken embryos and regulate valve progenitor cell formation (Barron et al., 2000; Lincoln et al., 2006; Zhang et al., 2010).

To our knowledge, there are few studies reporting the generation of the valve endothelial-like cells or VEPs from hPSCs (Neri et al., 2019). Although a clear definition and the molecular signature of the transient VEP cell types have been unavailable, it is generally agreed that the VEPs have some unique functional features, including the ability to undergo EndoMT, having higher proliferative and clonogenic potential compared to other ECs, and bearing a unique gene expression profile (expressing *MSX1*, *NFATC1*, *SOX9*, *JAG1/2*, *NOTCH1/3, HEY1* and *TBX2*) (Wu et al., 2013; Neri et al., 2019). The role of TGF-β, BMP and Notch signaling in cardiac cushion and valve formation has been well discussed (Garside e tal., 2013; Kruithof et al., 2012; Timmerman et al., 2017). Despite the great progress made in the field, however, there are a number of gaps of valve endocardial cell formation. For instance, how the valve endocardial cells (or VEPs) are defined and specified within the endocardium? which signaling pathways are involved in the process and what are the key downstream targets or mediators? how are the cardiac cushions (endocardial cushion cells) maintained after its formation? What controls the valve endocardial cell fate and how the EndoMT is initiated? How the heterogeneity of VECs is maintained during valve remodeling and maturation? what are the phenotypical difference between the AVC or OFT endocardial cells (VEPs) compared to the neighboring endocardial cells at a global or single cell levels? Answering these questions will undoubtedly deepen our understanding of valvulogenesis, and benefit the treatment of valve diseases. HPSC-based valvular cell differentiation that faithfully recapitulates the developmental program of in vivo valvulogenesis therefore provides a unique model to address these questions. In this work, using a combination of BMP4 and TGFb inferred from developmental insights of different animal models, we were able to generate the VEC-like progentior cells from hPSCs that resemble the primary VECs isolated from healthy human aortic valves. By analyzing the gene expression profile during hPSC differentiation into VEPs, we had some interesting and important observations/thoughts which allow us to tackle some of the above questions. First, we found that the crosstalk or synergy of BMP and TGF-β signaling pathways is important and sufficient for the specification of endocardial cells to a VEP fate (Lincoln et al., 2006). The requirement of BMP and TGF-β signaling pathways for valve endocardium formation has been described using animal models (Garside et al., 2013; Kruithof et al., 2012). However, whether this is true in human VEC formation remains unclear. We demonstrated that this mechanism might be conserved using the hPSC-based model. Second, we precisely dissected the signaling requirement during the VEP cell fate specification. In particular, we found that Notch signaling may not be strictly required during the initial stage of the VEP induction, although administration of DLL4 could enhance the expression of *NFATc1* (Supplementary Figure 2C). In stead, Notch signaling may be more important at later stage of VEP induction and its activation is required for the EndoMT process (Combs and Yutzey, 2012). This is in line with that Notch ligand (except *Notch1*) and receptor mutant mice appear largely normal before the onset of valve formation (Garside et al., 2013). Third, we found that VEP formation requires the activation of a cohort of target genes, in a time-dependent manner. It appears that *NFATc1* and *ETV2* are immediate early downstream targets, and Notch-related genes *HEY1/JAG1* are targets at later stage. As the Notch-related genes, including Notch ligands and its downstream genes *HAND2/SNAI1/TBX2/3,* are induced along with other VEGs such as *GATA4*, *NFATc1*, *TEK*, *EMCN* and *TGFβ1/2*, it is strongly suggested that Notch activation might be a part of VEP induction (Figure 3C). Fourth, hPSC-derived VEPs are unstable and prone to undergo EndoMT. We found that VEP derivation is associated with up-regulation of Notch-related genes *HEY1/JAG1*, bone formation-related genes and EndoMT factors SNAI1/SLUG/TWIST (Figure 3C; Supplementary Figure 3A, B). It is well-established that Notch downstream of TGF/BMP signaling may signal through SNAIl/SLUG to suppress *CDH5* expression in a subset of valve endocardial cells and facilitate the EndoMT (Garside et al., 2013; Combs and Yutzey, 2009; Wu et al., 2011), and/or through HEY1, JAG1 and GATA4, key mediators of EMT that are induced by TGF-β signaling (Yang et al., 2017; Zavadil et al., 2004).

An interesting observation in our current protocol is that constant BMP activities are needed throughout hPSC differentiation into VEPs (Figure 6A). This is not unexpected as BMP is one of the most important morphogens patterning early germ layers and cardiogenic mesoderm during embryogenesis (Kruithof et al., 2012; Jiao et al., 2003). Bmp4 is expressed in E8.0 when AVC is formed and is required for normal septation of the OFT (Jiao et al., 2003; liu et al., 2004). BMP expression in the AVC myocardium is necessary and sufficient to induce cushion formation and EMT of the adjacent endocardium (Palencia-Desai et al., 2015; Sahara et al., 2014; Sriram et al., 2015; Vincent and Buckingham 2010), and attenuating Bmp signaling inhibits SHF differentiation and the subsequent endocardial cushion formation (Brade et al., 2013; Combs and Yutzey, 2016; Hinton and Yutzey, 2011; Kattman, 2011; Lee et al., 1994; Lopez-Sanchez et al, 2015; McCulley et al., 2008; Nakano et al., 2016; Palpant et al., 2015; Rochais et al., 2009). BMP target genes *TGFb2, TBX2* and *TBX20* are known to be expressed in human valve endocardial cells and important for the endocardial cushion formation (Combs and Yutzey, 2009; Shelton and Yutzey, 2007; Singh et al., 2011). Importantly, fate mapping experiments showed that endocardial progenitor cells were restricted to the most ventral region of the embryos where high BMP activities exist (Lee et al., 1994). Mechanistically, BMP signaling may promote the earliest cardiac markers *NKX2.5* and *GATA4* as well as the earliest endothelial markers *Etv2* and *NFATc1* (Bai et al., 2010; Row et al., 2018; Zhou et al., 2004; Wu et al., 2011). Interestingly, we found that BMP ligand genes *BMP2/4/6* and BMP target *ID2* were abundantly expressed in both hPSC-derived VEPs (embryonic form) and the primary hVECs (adult form). Thus, we propose that BMP signaling is likely critical not only for embryonic valve development but also for adult valve homeostasis and function.

One major object using PSCs for regenerative medicine applications is the obtaining and culture of homogeneous populations of cells that are functional while retaining the potential for self-renewal and incorporation into matrices and tissues. To obtain the VEPs with sufficient purity and yield from hPSCs, it will be very useful if the VEC-specific markers are available. In this work, by performing comparative transcriptome analysis of HUVEC, HAEC and the freshly isolated VECs of different ages, we identified markers that are highly expressed in hPSC-derived VEPs and the primary VECs but lowly expressed in HUVEC and HAEC. In addition to the known VEGs such as *JAG1*, *HEY2*, *Notch3* and *DLL4*, some new candidate markers were identified, such as *CXCL12*, *HAND2*, *CDH11*, *PDGFRa*, *ISLR*, *LDB2*, *GLDN5*, *GYPC*, *WNT9b*, *EDN1* and *HMCN1* (Figure 4F). Some of these genes have been shown to be implicated with valve endocardial cell formation. For instance, endocardium expressed HAND2 was recently shown to control endocardium development via regulation VEGF signaling (VanDusen et al., 2015). The chemokine CXCL12 and its receptor CXCR4 are localized to the endocardial cells of the OFT and atrioventricular cushions, and disruption of this signaling causes cardiac defects (Sierro et al., 2007). CDH11 and PDGFRa has been shown to be expressed in valve progenitor cells (El-Rass et al., 2017). Surprisingly, we found that both hPSC-derived VEPs and the primary VECs expressed genes that are known to be associated with bone formation, antioxidative activities, shear stress and atherosclerosis (Figure 4E; Supplementary Figure 4F, G). This observation implied that hVECs might be actively involved in the progressive calcification and sclerosis of valves, which may explain why valve disease is closely linked with sclerotic and calcific lesions, and inflammation. In fact, signaling pathways and factors regulating valve development could be also involved in valve disease and congenital heart defects (Garside et al., 2013). Remarkably, KEGG analysis showed that hVECs are enriched with genes in TGF-β, PI3K/AKT, FGF, BMP and Notch signaling pathways, compared to HAEC/HUVEC (Figure 4E). In future, it will be important to investigate how these signaling crosstalk and converge to control valvulogenesis and maintain valve endothelial cells in physiological conditions.

HPSC-derived VEPs may represent an immature form of VEC-like cells, as the expression levels of shear stress response genes such as *KLF2*, *CAV1*, *NOS3* and *ICAM1* were significantly lower in hPSC-derived VEPs than the primary VECs (Supplementary Figure 4H). This was likely because hPSC-derived VEPs lack of mechanical stimulus that are exposed to hVECs in normal physiological conditions. A major limitation of this study is lack of comparative transcriptome analysis for hPSC-derived VEPs and embryonic VECs, due to the embryo inaccessibility and/or difficulties of obtaining the embryonic VECs. Thus, the precise maturity of hPSC-derived VEPs remains unclear. This is important when hPSC-derived VEPs are used for seed cells for the next generation TEVs. It has been shown that EC identity could be better maintained under continuous stimulation with shear stress or cyclic strain by using bio-reactors in vitro or by grafting the vessel into a host organism in vivo (Nejad et al., 2016). This suggested that a better strategy to obtain mature VEC-like cells is to stimulate the immature hPSC-derived VEPs by mechanical forces (such as bio-reactor) and tissue-engineered materials that can provide spatial and temporal control over the presentation of proper signaling molecules (cytokines or small molecules). We are currently working in this direction in the lab.

## Acknowledgement

We thank Prof. Donghui Zhang from Hubei University (Wuhan, China) for providing the PGP1 iPSC for us. We thank Prof. Ning Wang from Illinois University and Dr. Weihua Qiao from UNION hospital of HUST for the comments of the manuscript. This work was supported by National Key Research and Development Program of China (2016YFA0101100), National Natural Science Foundation of China (31671526) and Hundred Talents Program to YH Sun, and by National Key Research and Development Program of China (2016YFA0101100) and National Natural Science Foundation of China (81930052) to NG Dong.

## Materials and Methods

### Isolation of human VECs and HUVEC

Side-specific human aortic VECs [from the aortic side (VEC-A) and the ventricular side (VEC-V)] were isolated from non-calcified AVs obtained from heart transplant surgeries (according to an Institutional Review Board approved protocol at UNION hospital, HUST) using a brief collagenase digestion and gentle scraping method as previously described (Holliday et al., 2011). Briefly, Human aortic valves from donors with no history of cardiovascular medical conditions were obtained postmortem. Human aortic VECs from 7-, 9-, 19-, 30-, 40-, 50- and 59-year old were isolated and cultured with ECGM or EBM-2 medium (Lonza) for 1-2 days to examine the cell morphology. To keep the intact features of the isolated VECs, the cells were not passaged and were directly used for RNA-sequencing. The quality of isolated hVECs was evaluated by three criteria. First, the primary VECs were checked morphologically under the microscope. The batch of the isolated VECs with significant contamination with fibroblast-like cells (likely the VICs) was not used for RNA-sequencing. The vast majority of the primary VECs (> 99%) adopted a cobblestone morphology, suggesting that the isolated VECs were not significantly contaminated by the VICs. Second, to further reduce the possibility that the isolated VECs were contaminated by the VICs or other non-valvular cell types, cells with fibroblastic morphology were manually removed. Third, the RNA-seq data were used to double-check the quality of the isolated primary VECs. The RNA-seq results showed that the definitive EC markers *CDH5* and *PECAM1* were highly expressed while the fibroblast markers *S100A4* and *POSTN* as well as cardiomyocyte markers *TNNT1/2* were barely detectable (RPKM< 5; *P*< 0.05).

Human endothelial cell line HUVEC (#CRL-1730) was purchased from the American Type Culture Collection (Rockville, MD, USA). Human aortic endothelial cell line HAEC (#6100) was purchased from Shanghai Cell Bank, CAS (Shanghai, China).

### Maintenance of human PSCs

Human PSCs including human induced pluripotent stem cells (Colunga et al., 2019) and PGP1 human induced pluripotent stem cells (kindly provided by Prof. Zhang from Hubei University, Wuhan, China), and human ESC lines (H8 and H9 lines, Wicell, WI, USA) were used for this study.

HPSCs were maintained with mTsSR1 medium (STEMCELL Technologies, Canada) or E8 medium (Life Technologies, USA), in feeder-free plates coated with Matrigel (Corning) as previously described (Berger et al., 2016). HPSCs were treated with Accutase (Gibco) for 4 min, and passaged every four days (Singh et al., 2012; 2015). Briefly, single hPSCs were centrifuged at 400 g for 4 min, and seeded onto the Matrigel-coated 6-well plates at 30,000 cell/cm^2^ in mTeSR1 supplemented with 0.1 μM ROCK inhibitor Y-27632 (Tocris, USA).

### CPC differentiation

We have previously described the method for the generation of CPCs from hPSCs (Berger et al., 2016). To optimize the generation of ISL1^+^ KDR^+^ CPCs, we further modified the method. HPSCs were passaged as described above and reseeded at 5× 10^5^ cells/cm^2^ (H9) or 8× 10^4^ cells/cm^2^ (H8 and iPSCs) onto the Matrigel coated plates with Essential 6 medium (Gibco) supplemented with WNT3a (5036-WN, R&D Systems) and BMP4 (314-BP, R&D Systems) for 1 day. Next, cells were cultured in E6 medium supplemented with BMP4 and bFGF for 3 days with daily medium change.

### CPCs to endocardial precursor cells

hPSC-derived day 3 CPCs were dissociated and seeded at 1× 10^5^ cells/cm^2^ onto cell culture dishes. To generate endocardial progenitor cells, the seeded cells were cultured with Essential 6 medium supplemented with VEGF, bFGF and BMP4. We indeed found that the addition of 10-20 ng/ml of DLL4 increased the expression of *NFATc1* as discussed in the text. The medium was changed daily for 3 days.

### Endocardial precursor cells to the VEPs

HPSC-derived day 6 endocardial cells were treated with TrypLE solution (Gibco) for 4 min and passaged at the ratio of 1: 2, and were cultured for 3-9 days using E6 medium supplemented with BMP4 and TGFb. The medium was changed every two days.

### RNA-seq and heat map analysis

HAEC, HUVEC, H9 cells, HFF, the primary VICs, the primary VECs and hPSC-derived VEPs were collected and lysed with 1ml Trizol (Transgen Biotech, China). RNA sample quality was checked by the OD 260/280 value using the Nanodrop 2000 instrument. When necessary, hPSC-derived VEPs were sorted with VE-cad or CD31 magnetic beads (Miltenyi Biotec, GmbH, Germany) before RNA-sequencing. RNA samples were sent to the BGI China where RNA-sequencing libraries were constructed and sequenced by a BGI-500 system. RNA-seq experiments were repeated at least 2 times. Differentially expressed genes (DEGs) were defined by FDR < 0.05 and a Log2 fold change >1 was deemed to be DEGs. The heat map was constructed based on the commonly expressed genes in different cell types, and the top 75 DEGs were listed as part of the heap map. Gene ontology (GO) analysis for differentially expressed genes (DEGs) and heat maps were generated from averaged replicates using the command line version of deepTools2.

### Quantitative real-time PCR

Total RNA for cells was extracted with a Total RNA isolation kit (Omega, USA). 1 μg RNA was reverse transcribed into cDNA with TransScript All-in-One First-Strand cDNA synthesis Supermix (Transgen Biotech, China). Quantitative real-time PCR (RT-qPCR) was performed on a Bio-Rad qPCR instrument using Hieff qPCR SYBR Green Master Mix (Yeasen, China). The primers used for RT-qPCR are listed in Table 2. All experiments were repeated for three times. The relative gene expression levels were calculated based on the 2^-ΔΔCt^ method. Data are shown as means ± S.D. The Student’s t test was used for the statistical analysis. The significance is indicated as follows: *, p < 0.05; **, p < 0.01; ***, p < 0.001.

**Table 2.**
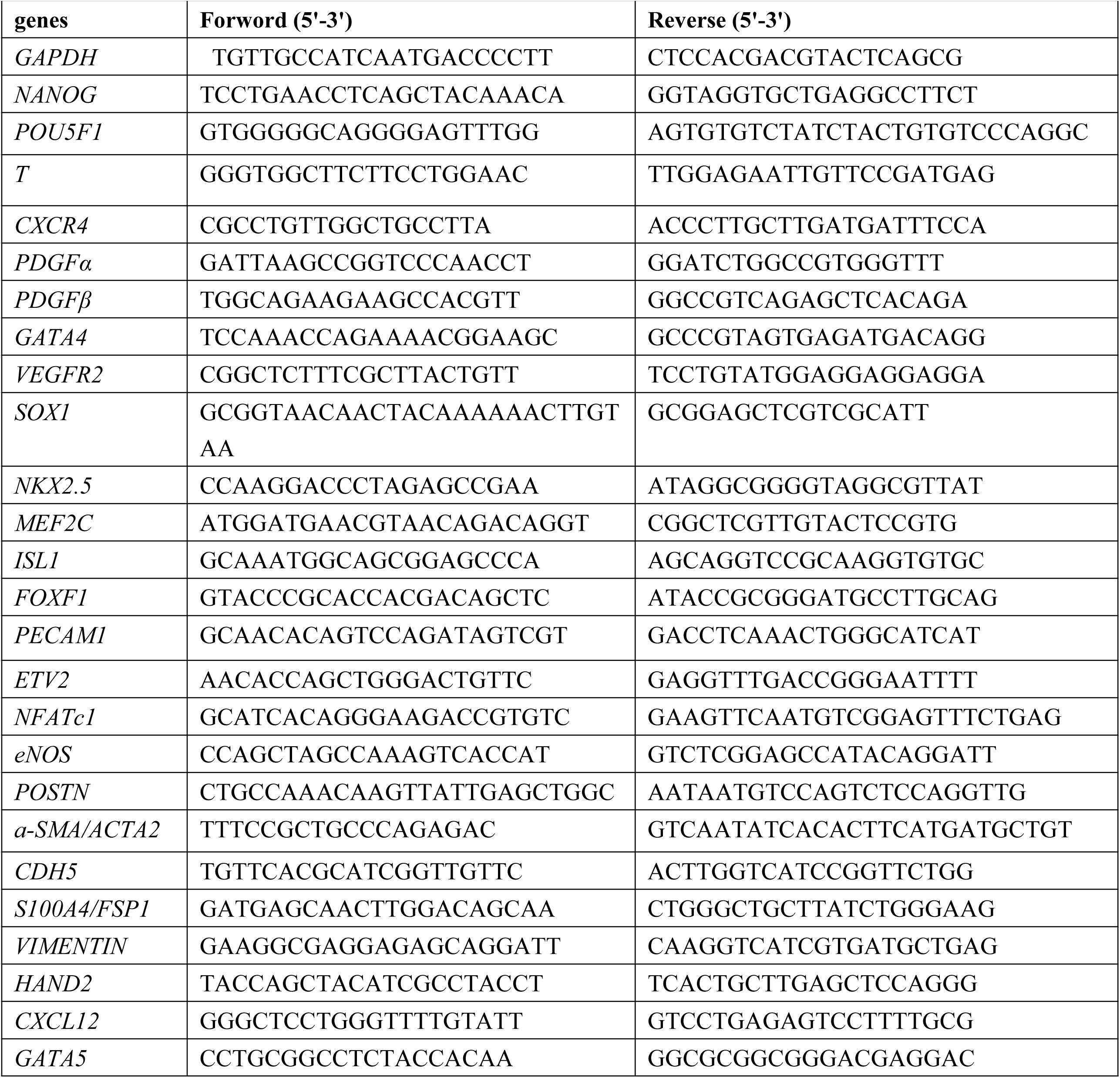
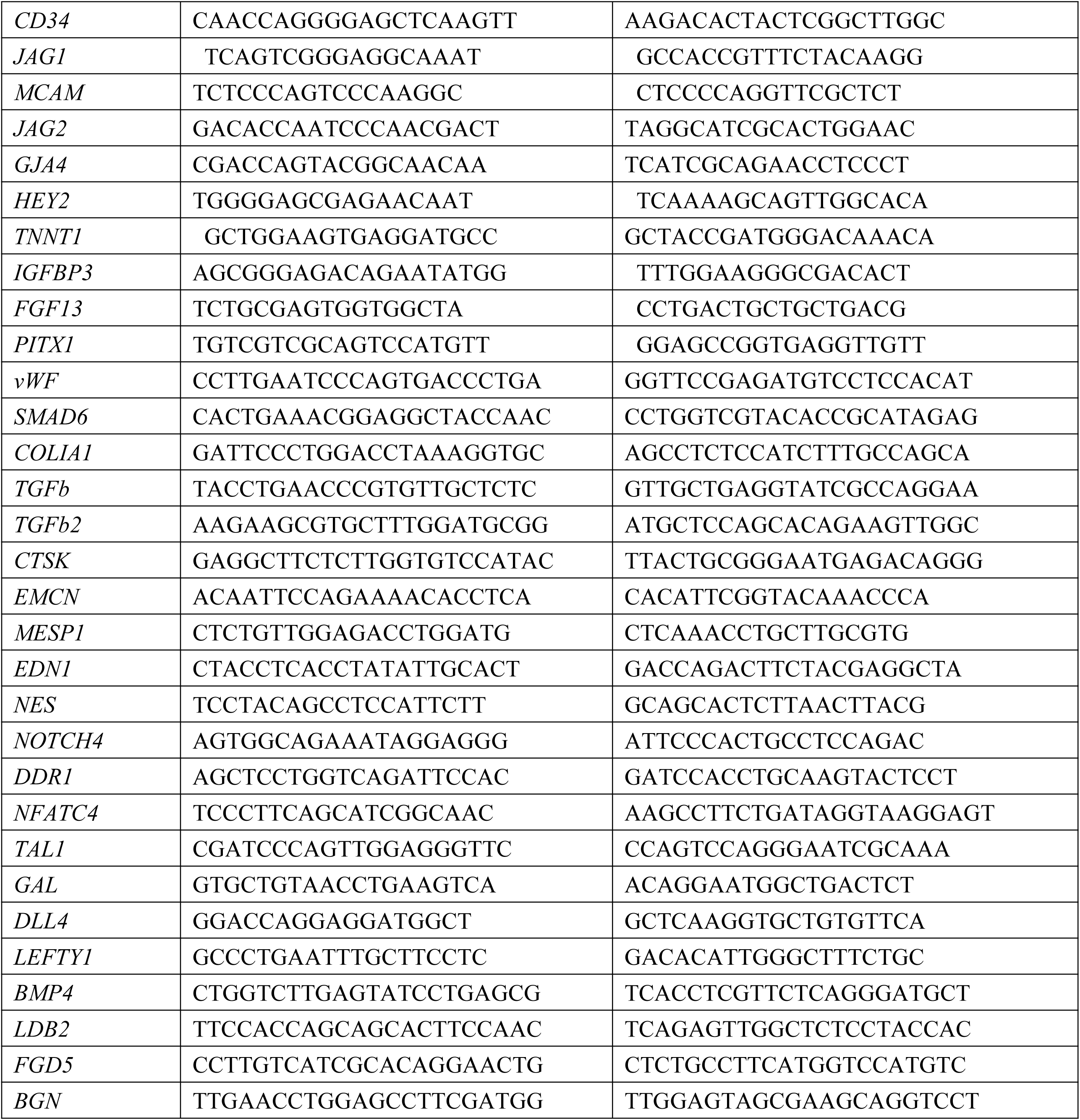
The primers used for qRT-PCR analysis genes Forword (5’-3’) Reverse (5’-3’)

### Western blot analysis

WB analysis was done as previously described (Berger et al., 2016). Briefly, cells were lysed on ice in SDS lysis buffer (50 mM Tris-HCl, 150 mM NaCl, 5 mM EDTA, 1% TritonX-100, 0.5% Na-Deoxychote, 1× Protein inhibitor, 1× DTT) for 30 min, shaking for 30s every 5 min. Protein samples were resolved by SDS-PAGE (EpiZyme) and transformed to the PVDF membranes. The blots were incubated over night at 4℃ with primary antibodies against Anti-CD31 (Abcam, ab28364, 1:500), Anti VE-cadherin (Abcam, ab33168, 1:1000), Anti-NFATc1 (Abcam, ab2796, 1:2000), Anti-ISLET1 (Abcam, ab20670, 1:1000), FSP1 (Abcam, ab124805, 1:1000), anti-TBX2 (Proteintech, 22346-1-AP, 1: 200), Anti-BMP4 (Abcam, ab39973, 1:1000) and Anti-GAPDH (Santa Cruz, sc-25778, 1:1000), followed by incubation with a HRP-conjugated goat anti-rabbit IgG (GtxRb-003-DHRPX, ImmunoReagents, 1:5000), a HRP-linked anti-mouse IgG (7076S, Cell Signaling Technology, 1:5000) for 1 hour at room temperature. Western blotting was detected by ECL substrate (Advansta, K-12045-D20) and visualized by LAS4000 mini luminescent image analyzer (GE Healthcare Life Sciences, USA).

### Flow cytometry assay

Cells were washed twice with DPBS (BI, China), and digested with Typsin (BI) for 1 min, folowed by wash with DPBS containing 0.5% BSA (0.5% PBSA). Next, cells were centrifuged and washed once with 0.5% PBSA. The resuspended cells in 200 ul 0.5% PBSA were passed the flow tube to obtain single cells. After that, cells were incubated with the antibodies for 30 min at room temperature. The antibodies used are Anti-NFATc1 (Abcam, ab2796); Anti-ISLET1 (Abcam, ab178400); Anti-VE-cadherin (Abcam, ab33168), Anti-CD31 (R&D, FAB35679), Anti-KDR (R&D, FAB3579), Anti-CDH5 (BD Horizon, 561569), Anti-SOX9 (Proteintech, 67439-1-Ig, 1: 200), Anti-HEY1 (Proteintech, 19929-1-AP, 1: 200), Anti-JAG1 (Proteintech, 668909-1-Ig, 1: 200) or (Invitrogen, PA5-86057), Anti-NOTCH1 (Proteintech, 20687-1-AP, 1: 200), Anti-P-SELECTIN (Proteintech, 60322-1-Ig, 1: 200), Anti-PROX1 (Proteintech, 11067-2-AP, 1: 200) and the isotype control antibodies: mouse IgG (R&D, C002P) and rabbit IgG (Invitrogen, 10500C). After 3 times washing, cells were treated with secondary antibodies at room temperature for 15 min. Finally, cells were washed twice and resuspended with 300-500 μl 0.5% PBSA, and then analyzed by Accuri C6 flow cytometer (BD Biosciences, USA).

### Immunofluorescence

Cells were fixed in 4% paraformaldehyde (PFA) for 15 min at room temperature, then were washed 3 times with DPBS containing 5% Triton X-100 for 10 min. Following the incubation with blocking buffer (5% normal horse serum, 0.1% Triton X-100, in PBS) at room temperature for 1 hour, cells were incubated with primary antibodies at 4℃ overnight. The primary antibodies used were: ISL1 (Abcam, ab178400, 1:300), CD31 (Abcam, ab28364, 1:80), NOTCH4 (Abcam, ab225329, 1:100), VE-cadherin (Abcam, ab33168, 1:300), CDH5 (proteintech, 66804-4-lg, 1:200), NFATc1 (Abcam, ab2796, 1:100), Endomucin (Abcam, ab106100, 1:50), HEY1 (Proteintech, 19929-1-AP, 1: 200), LDB2 (Abcam, ab3627, 1:100), GATA4 (Proteintech, 19530-1-AP, 1:100) and NESTIN (Abcam, ab22035, 1:200). After three-times washing with PBST, the cells were incubated with secondary antibodies (1: 500 dilution in antibody buffer, Alexa Fluor-488 or -555, ThermoFisher) at room temperature for 1 hour in the dark. The nuclei were stained with DAPI (D9542, Sigma, 1:1000). After washing with PBS twice, the slides were mounted with 100% glycerol on histological slides. Images were taken by a Leica SP8 laser scanning confocal microscope (Wetzlar, Germany).

Quantification of immunofluorescence staining was done by ImageJ software. When measuring the number of positive cells, 3-4 random fields per coverslip were counted. DAPI-positive cells (a total of appropriately 500 cells) were counted as the total number of cells. The proportion of cells positive for specific markers was calculated with respect to the total number of DAPI-positive cells, and the results were expressed as the mean ± s.e.m. of cells in 5-6 fields taken from 3 to 4 cultures of three independent experiments. Differences in means were statistically significant when *p*< 0.05. Significant levels are: \**p*< 0.05; \*\**P*< 0.01.

### Chromatin Immunoprecipitation (ChIP)

ChIP experiments were performed according to the Agilent Mammalian ChIP-on-chip manual as described (Singh et al., 2012). Briefly, 1×10^8^ cells were fixed with 1% formaldehyde for 10 min at room temperature. Then the reactions were stopped by 0.125 M Glycine for 5 min with rotating. The fixed chromatin were sonicated to an average of 500-1,000bp (for ChIP-qPCR) using the S2 Covaris Sonication System (USA) according to the manual. Then Triton X-100 was added to the sonicated chromatin solutions to a final concentration of 0.1%. After centrifugation, 50 μl of supernatants were saved as input. The remainder of the chromatin solution was incubated with Dynabeads previously coupled with 10 μg ChIP grade antibodies (SMAD4, #9515, CST) overnight at 4℃ with rotation. Next day, after 7 times washing with the wash buffer, the complexes were reverse cross-linked overnight at 65℃. DNAs were extracted by hydroxybenzene-chloroform-isoamyl alcohol and purified by a Phase Lock Gel (Tiangen, China). The ChIPed DNA were dissolved in 100 μl distilled water. Quantitative real-time PCR (qRT-PCR) was performed using a Bio-Rad qPCR instrument. The enrichment was calculated relative to the amount of input as described. All experiments were repeated at least for three times. The relative gene expression levels were calculated based on the 2^- ΔΔCt^ method. Data were shown as means ± S.D. The Student’s t test was used for the statistical analysis. The significance is indicated as follows: *, *p* < 0.05; **, *p* < 0.01; ***, *p* < 0.001.

### Low-density lipoprotein uptake assay

HPSC-derived VEPs, HUVEC and H9 ESCs (as negative control) were cultured to a confluency of 30-40%. After serum starve of the cells for 12 hours, cells were incubated at a final concentration of 15 ng/ml of Alexa Fluor 594 AcLDL (Invitrogen) for 4 hours at 37℃. The cells were rinsed twice with DPBS and were fixed with 4% PFA for 10 min at room temperature. The nuclei were stained with DAPI (D9542, Sigma, 1:1000). Images were taken by a Leica SP8 laser scanning confocal microscope (Wetzlar, Germany).

### In vitro tube formation assay

HPSC-derived VEPs, HUVEC and the isolated VECs were cultured on a Matrigel (Corning) coated 96-well plate (Matrigel, #356234, Corning, USA) in EBM-2 medium (Lonza). Images were taken 24 hours later after plating under a phase-contrast microscope.

### Interaction between the de-cellularized porcine valves and the VEPs

De-cellularized porcine aortic valves were prepared as previously described (Hu et al., 2013; Zhou et al., 2013). Approximately 5×10^5^ hPSC-derived VEPs, the isolated primary VECs and the control HUVEC were added to the apical surface of the DCV constructs and allowing attachment for 24 hours in a CO_2_ incubator. Then the medium was removed and tissue constructs were carefully turned over using sterile forceps, and cells seeding procedure was repeated on the opposite surface of the constructs. The constructs were maintained in culture for a week with medium changed daily or 3-4 weeks to study the long term interaction between the seeded VEPs and the de-cellularized porcine aortic valves.

### Transwell assay

HPSC-derived VEPs and the isolated VECs were washed twice with PBS. A total of 8× 10^4^ cells were suspended in 200 μl serum-free EBM-2 and seeded in the upper chamber of a transwell system (3422, Corning, USA). The lower chamber was filled with 600 μl EBM-2 containing supplements and growth factors. Cells were allowed to migrate for 6 hours before membranes were fixed with 4% PFA for 10 min and stained with 0.5% crystal violet dyes for 2 hour. The cells on the upper chambers was scraped with a cotton swab. Random fields were photographed under a microscope and cells were counted.

### TUNEL staining assay

HPSC-derived VEPs and the isolated primary VECs were seeded onto a 96-well plate. After washing twice with PBS, cells were treated with H2O2 (200 μM) for 24h. Then TUNEL staining was performed using a TUNEL detection kit (Vazyme biotech Co. Ltd., China) according to the manufacturer’s instructions. The microscopic areas were randomly selected and the TUNEL-positive cells were calculated by Image-Pro Plus.

### EdU incorporation analysis

HPSC-derived VEPs, the isolated primary VECs and HUVEC were seeded on the de-cellularized porcine aortic valves. The EBM-2 culture medium and our home made medium (VEGFA+ BMP4+ bFGF) was added with 5-ethynyl-2-deoxyuridine (EdU, Ribobio, China). Cells were fixed at 72h and 96h after plating and processed for immunofluorescence by using the Cell-Light EdU Apollo567 In Vitro Kit (C10310-1, Ribobio, China). Images were captured and the number of EdU positive nuclei was counted manually by using Image-Pro Plus software.

### Cell viability analysis

HPSC-derived VEPs, the isolated VECs and HUVEC were seeded the de-cellularized porcine aortic valves, and seeded at the density of 7× 10^3^ cells/well. At day 1, 3, 5, and 7 after plating, cells were rinsed with PBS and cultured in 100 μl EBM-2, followed by the addition of 20 μl MTS solution (Promega, USA). A triple number of wells were set for each time point. Cell viability was measured with a spectrophotometer at an absorbance of 490 nm after 1h incubation.

### Data availability

All RNA-seq data have been deposited into the database at https://bigd.big.ac.cn/. The accession number is PRJCA002549. All other related data will be available upon reasonable request.

## Figure Legends

**Supplementary Figure 1.**
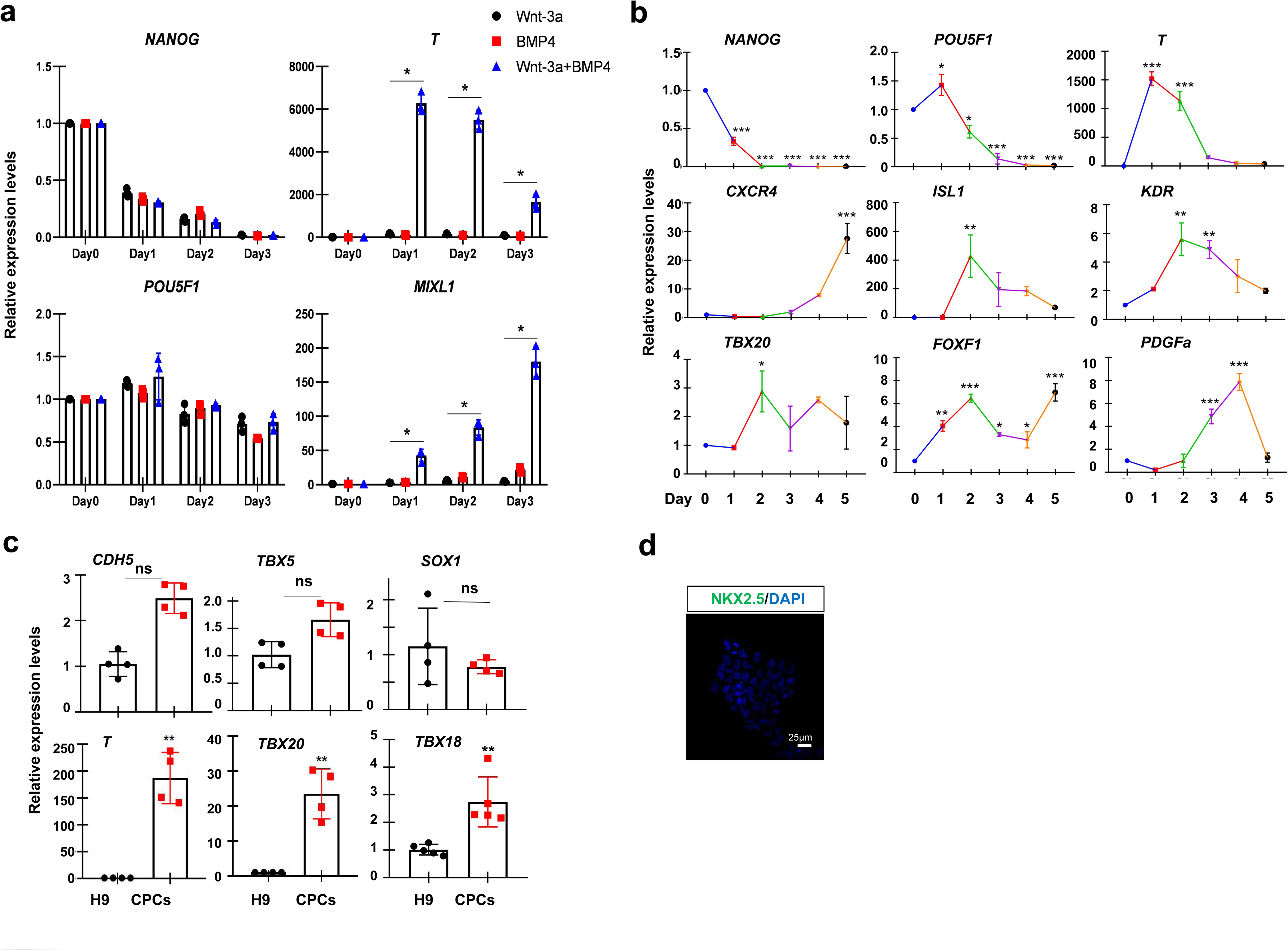
Related to Figure 1. **a** The qPCR analysis of T and MIXL1 for hPSCs treated with BMP4 and Wnt3a for a time course of 3 days. **b** Day 1 hPSC-derived T positive cells were treated with bFGF and BMP4 for a time course of 6 days, and dynamic expression of indicated genes were examined daily. **c** The qPCR analysis of hPSC-derived day 3 CPCs for the indicated markers. **d** IF staining of day 3 hPSC-derived CPCs showing the expression levels of NKX2.5. All experiments were repeated 3 times. Significant levels are: \**p*< 0.05; \*\**P*< 0.01; \*\*\**P*< 0.001. Shown are representative data.

**Supplementary Figure 2.**
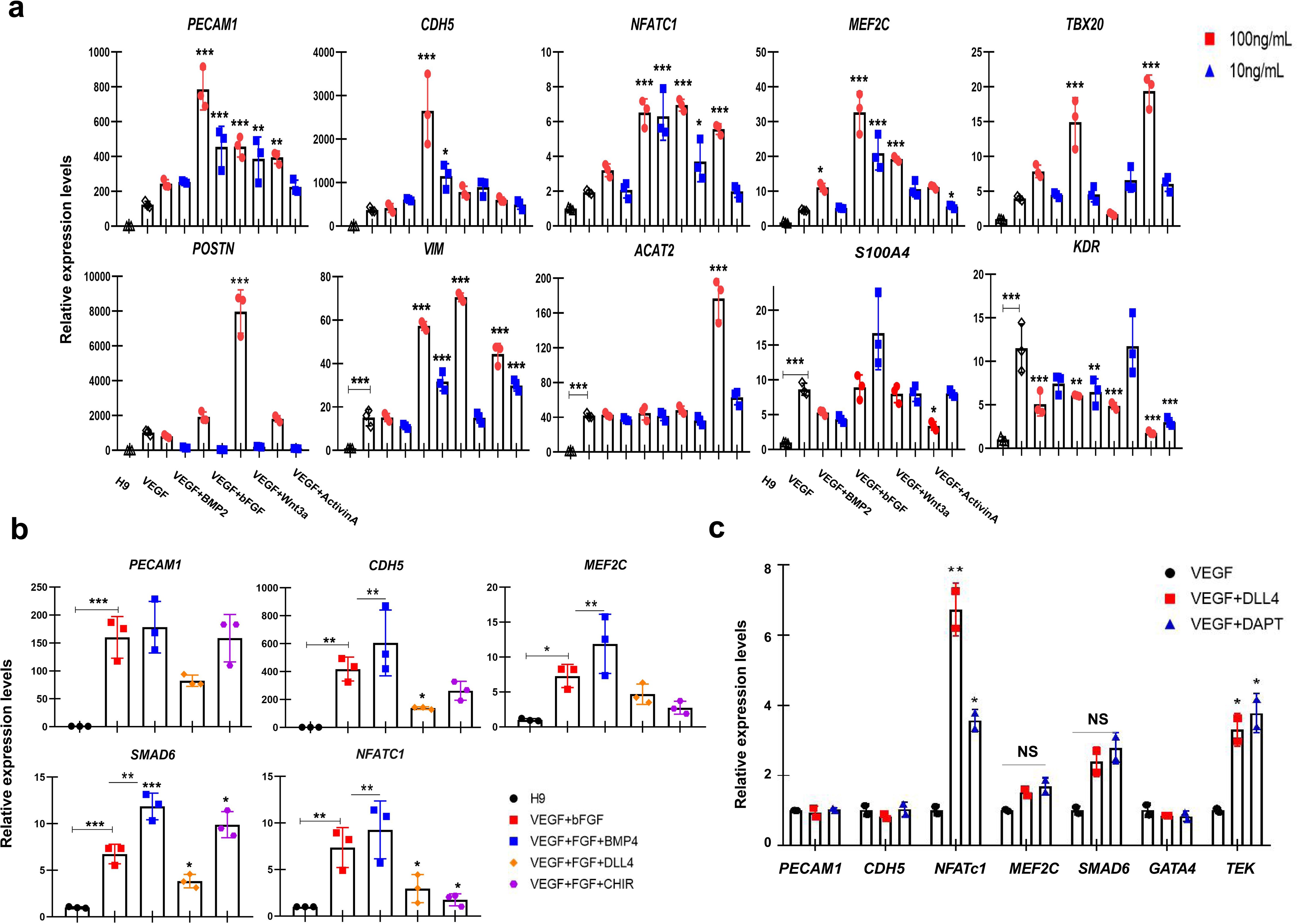
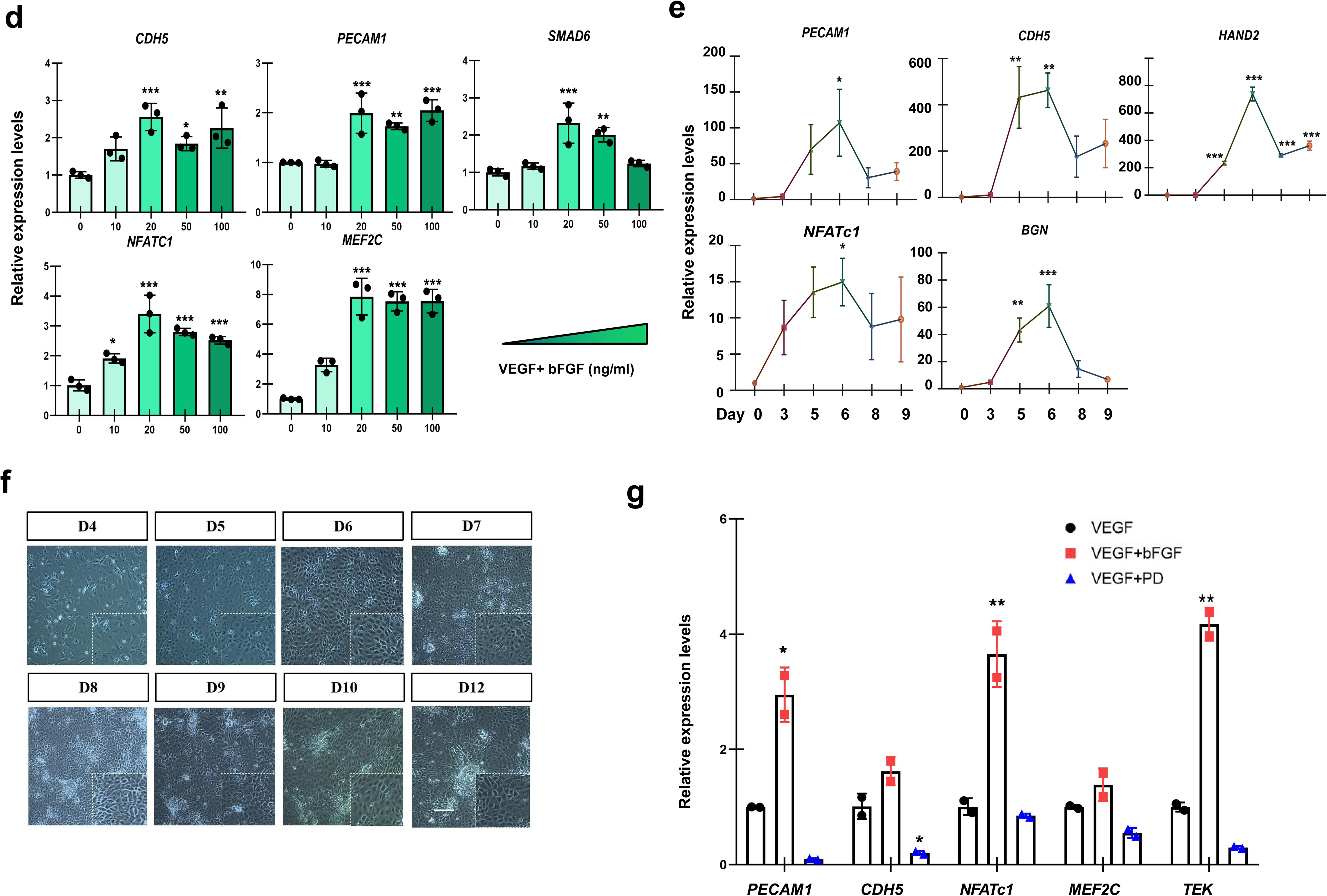
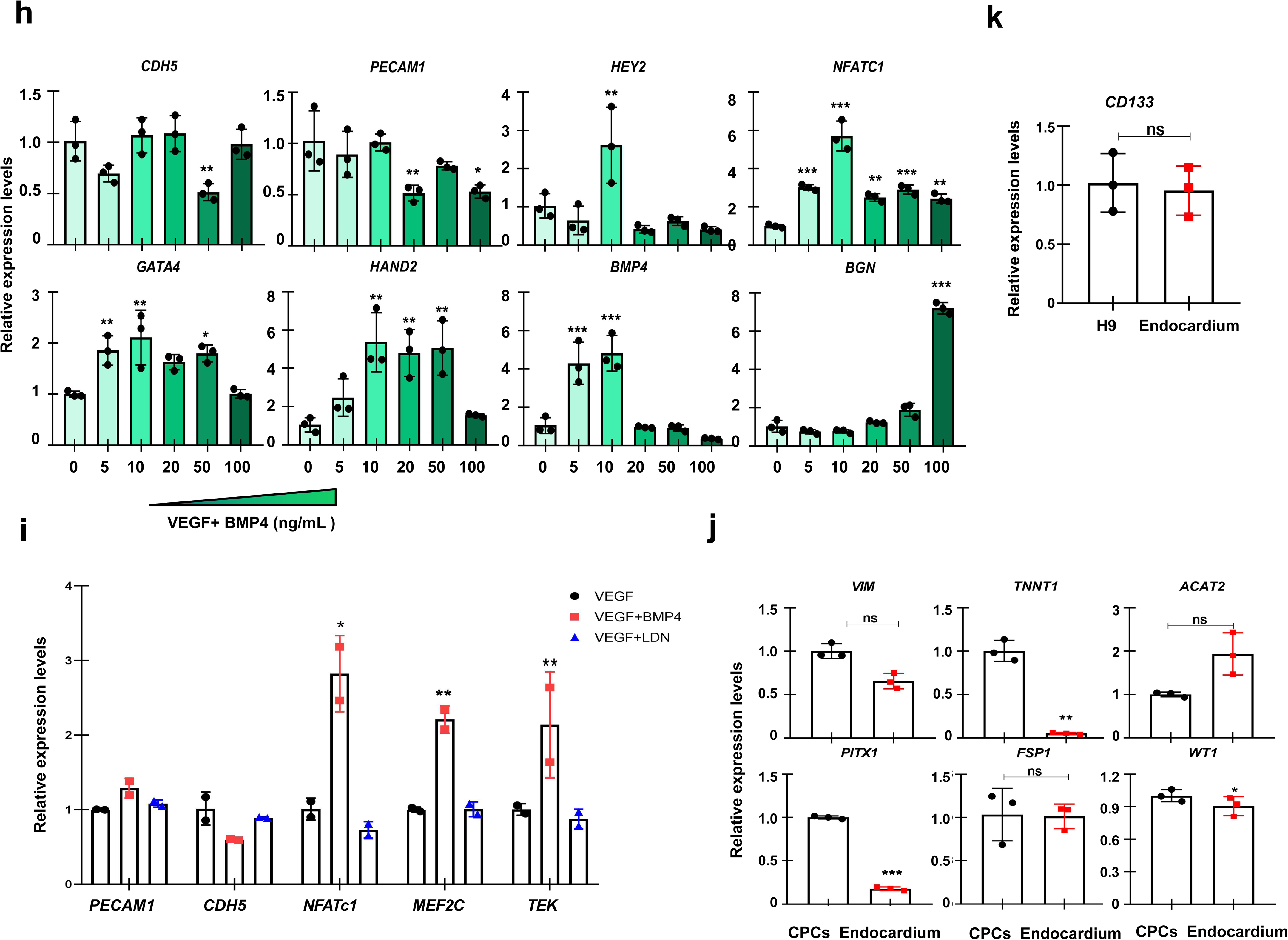
Related to Figure 2. **a** The qPCR analysis of the endocardium markers as well as VIC markers of day 3 hPSC-derived CPCs treated with the indicated signaling molecules with low and high concentration. **b** The qPCR analysis of the endocardium markers of day 3 hPSC-derived CPCs treated with the indicated signaling or small molecules. **c** The qPCR analysis of the endocardium markers of hPSC-derived CPCs treated with the Notch signaling activator DLL4 and inhibitor DAPT. **d** The qPCR analysis of the endocardium markers of hPSC-derived CPCs treated with an increasing doses of bFGF in the presence of VEGFA. **e** Dynamic expression analysis of the endocardium markers of day 3 hPSC-derived CPCs treated with VEGF and bFGF for a time course of 9 days. **f** The morphology of day 3 hPSC-derived CPCs treated with VEGF and bFGF for a time course of 12 days. **g** The qPCR analysis of the endocardium markers of hPSC-derived CPCs treated with FGF signaling ligand bFGF and its inhibitor PD. **h** The qPCR analysis of the endocardium markers of hPSC-derived CPCs treated with increasing dose of BMP4 in the presence of VEGF. **i** The qPCR analysis of the endocardium markers of hPSC-derived CPCs treated with BMP signaling ligand BMP4 and its inhibitor LDN. **j** The qPCR analysis of the indicated markers of hPSC-derived CPCs treated with VEGF, bFGF and BMP4 for 3 days. **k** The qPCR analysis of *CD133* of hPSC-derived CPCs treated with VEGF, bFGF and BMP4 for 3 days. All experiments were repeated 3 times. Significant levels are: \**p*< 0.05; \*\**P*< 0.01; \*\*\**P*< 0.001. NS: not significant. Shown are representative data.

**Supplementary Figure 3.**
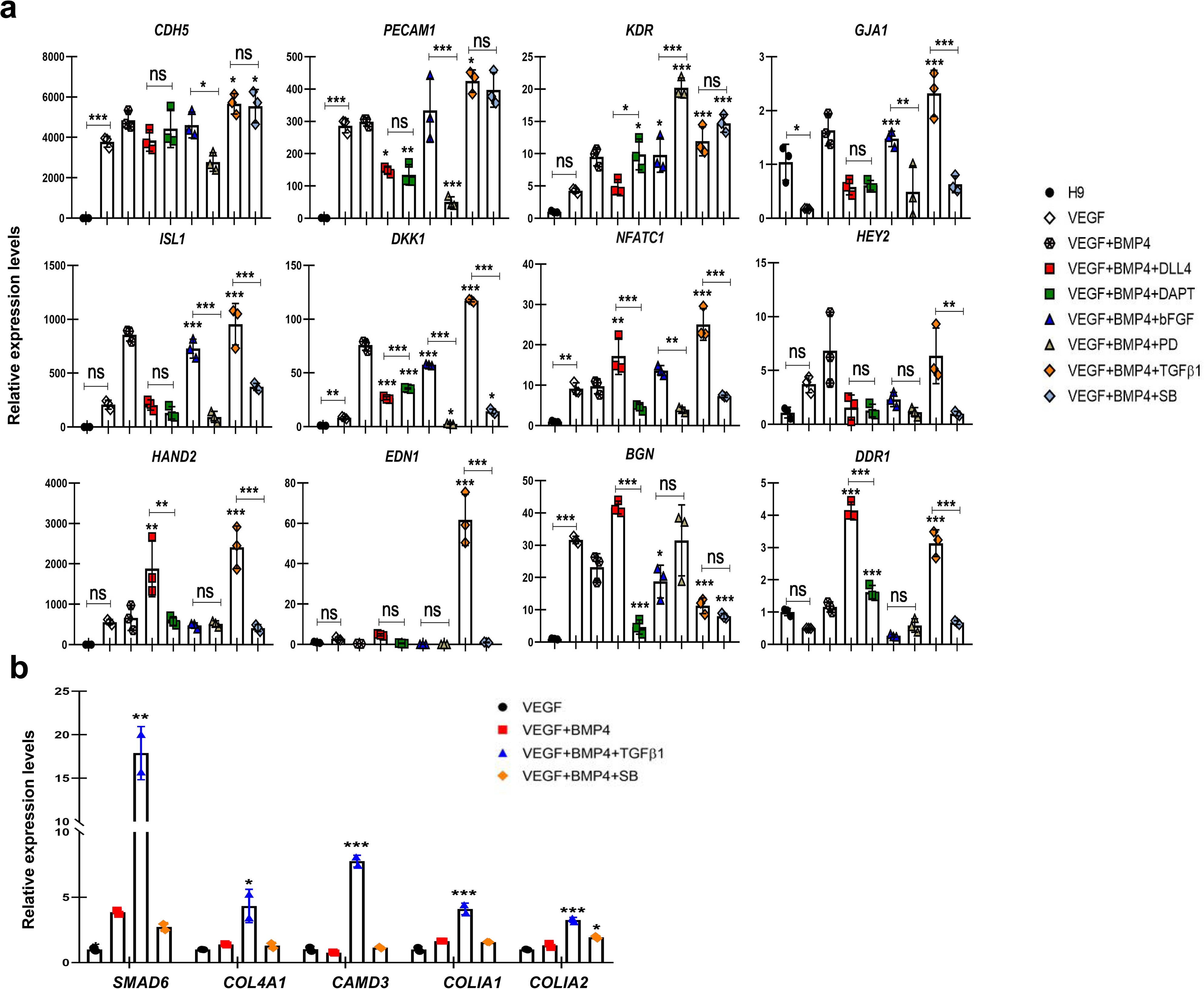
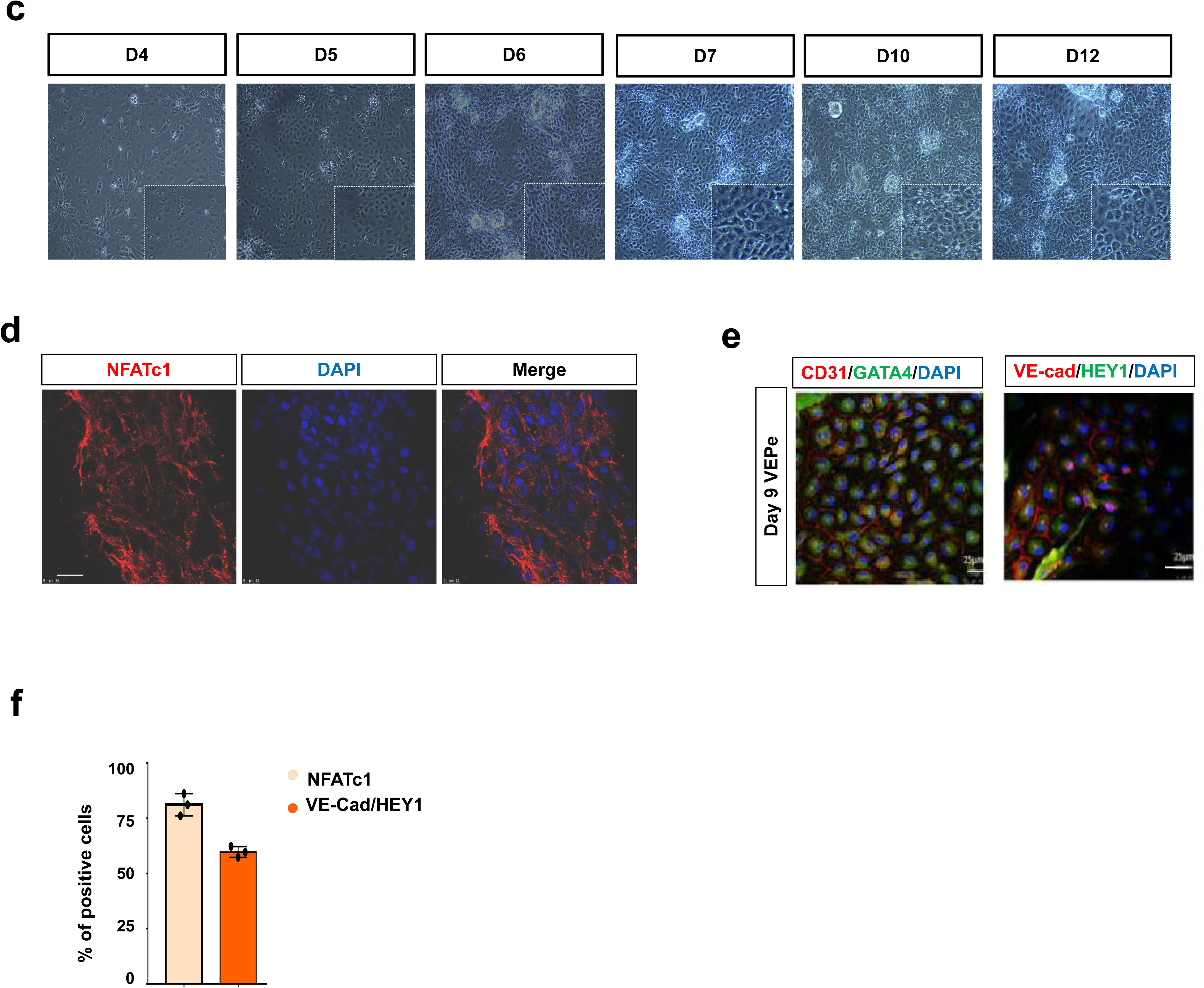
Related to Figure 3. **a** The qPCR analysis of the selected markers for day 9 hPSC-derived VEPs. Note that the combined treatment with VEGFA, BMP4 and TGFb leads to higher expression of the endocardium-enriched genes than other combined treatment. **b** The qPCR analysis of the indicated markers for day 9 hPSC-derived VEPs treated with TGF signaling ligand TGFb and its inhibitor SB. **c** The morphology of hPSC-derived CPCs treated with VEGF, TGFb and BMP4 for a time course of 12 days. **d** IF staining of day 9 hPSC-derived VEPs, showing the expression of NFATc1. **e** IF staining of day 9 hPSC-derived VEPs, showing the co-expression of CD31/GATA4 and VE-cad/HEY1. Scale bar: 25 μm. **f** Quantitative analysis of IF staining showing the percentage of positive cells for panels (d) and (e). All experiments were repeated 3 times. Significant levels are: \**p*< 0.05; \*\**P*< 0.01; \*\*\**P*< 0.001. Shown are representative data.

**Supplementary Figure 4.**
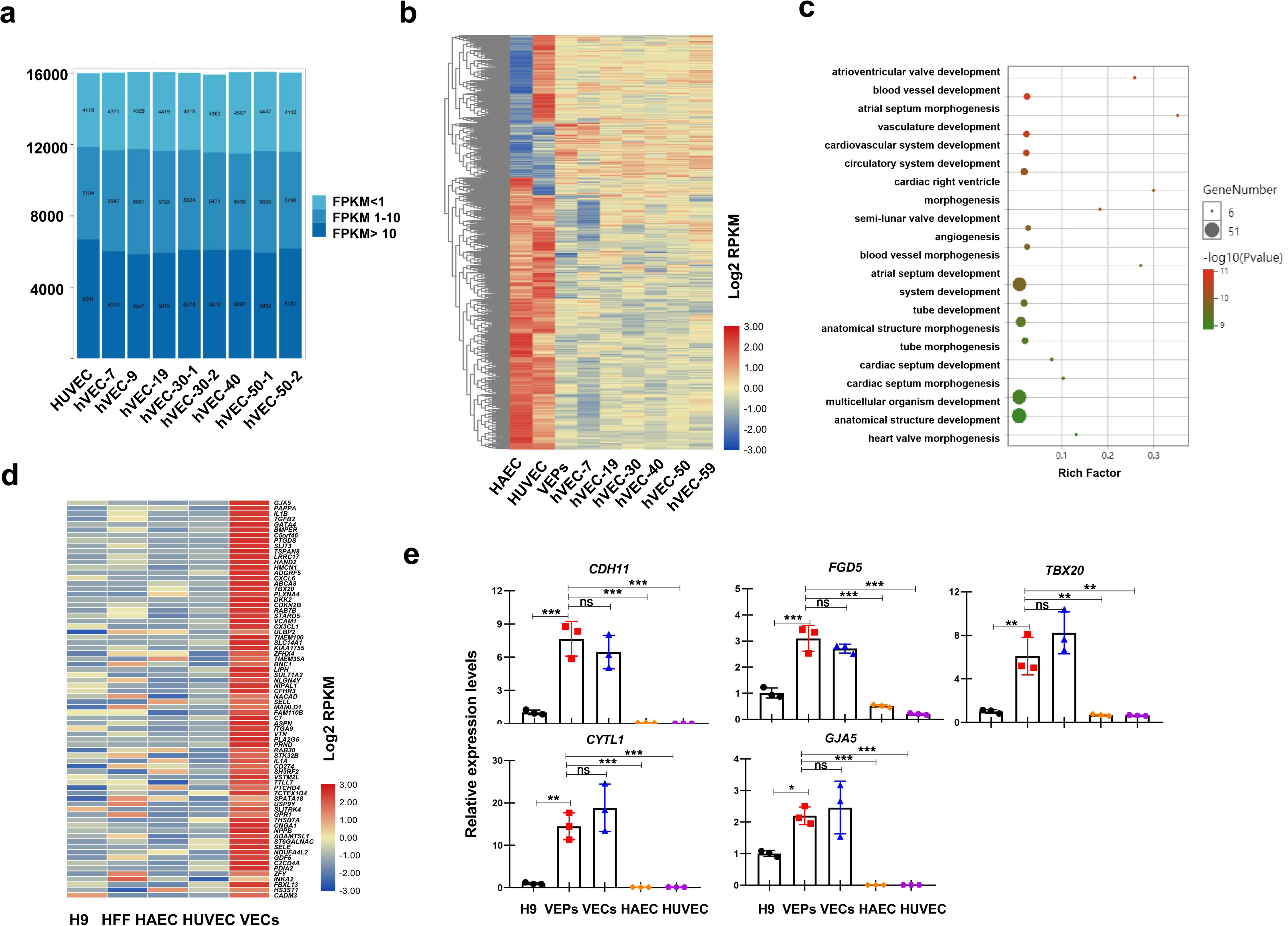
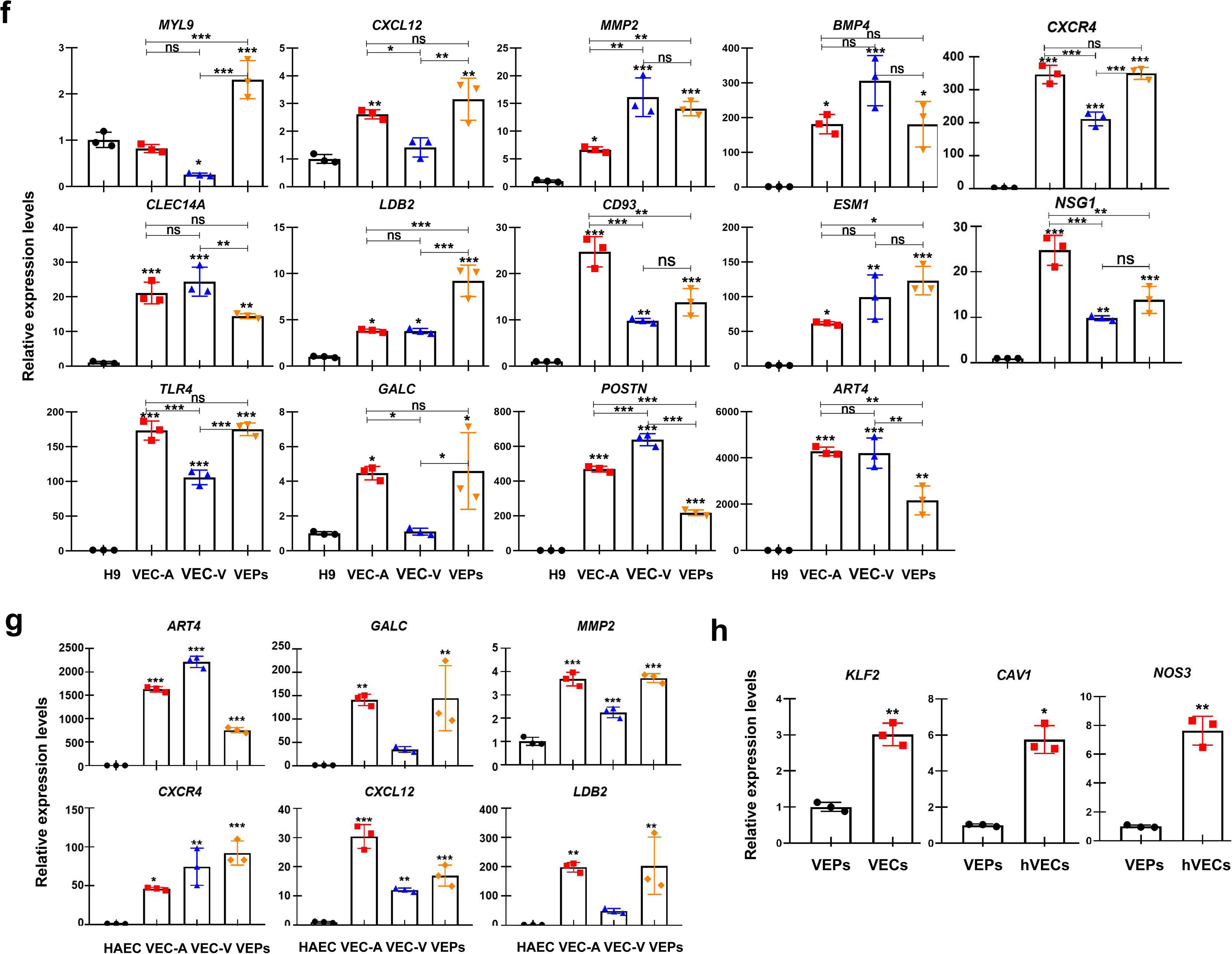
Related to Figure 4. **a** Representative image showing gene numbers and gene expression level of HUVEC and the primary VECs isolated from aortic valves of different ages. hVEC-30-1 and hVEC-30-2 are aortic valve endothelial cells from two 30-year-old individuals. **b** Heat map showing the relationship among HAEC, HUVEC, the primary VECs from different ages and day 9 hPSC-derived VEPs. Note that the expression profile of day 9 hPSC-derived VEPs is more similar to the primary VECs than HUVEC and HAEC. **c** GO result showing the enrichment of terms associated with cardiovascular and valve development. **d** Heat map showing top 75 genes that are highly expressed in hPSC-derived VEPs and the primary VECs but low in H9, HFF, HAEC and HUVEC. **e** The qPCR analysis of selected genes showing that they are highly expressed in both hPSC-derived VEPs and the primary VECs but lowly expressed in HUVEC and HAEC. **f** The qPCR analysis of a panel of markers showing that they are highly expressed in both day 9 hPSC-derived VEPs and the primary VECs but lowly expressed in H9. hVEC-A and hVEC-V stand for the primary VECs isolated from aortic side and ventricular side, respectively. **g** The qPCR analysis of a panel of markers showing that they are highly expressed in both day 9 hPSC-derived VEPs and the primary VECs but lowly expressed in HAEC. hVEC-A and hVEC-V are the primary VECs isolated from aortic side and ventricular side, respectively. **h** The qPCR analysis of shear stress response genes for day 9 hPSC-derived VEPs and the primary VECs. All experiments were repeated 3 times. Significant levels are: \**p*< 0.05; \*\**P*< 0.01; \*\*\**P*< 0.001. Shown are representative data.

**Supplementary Figure 5.**
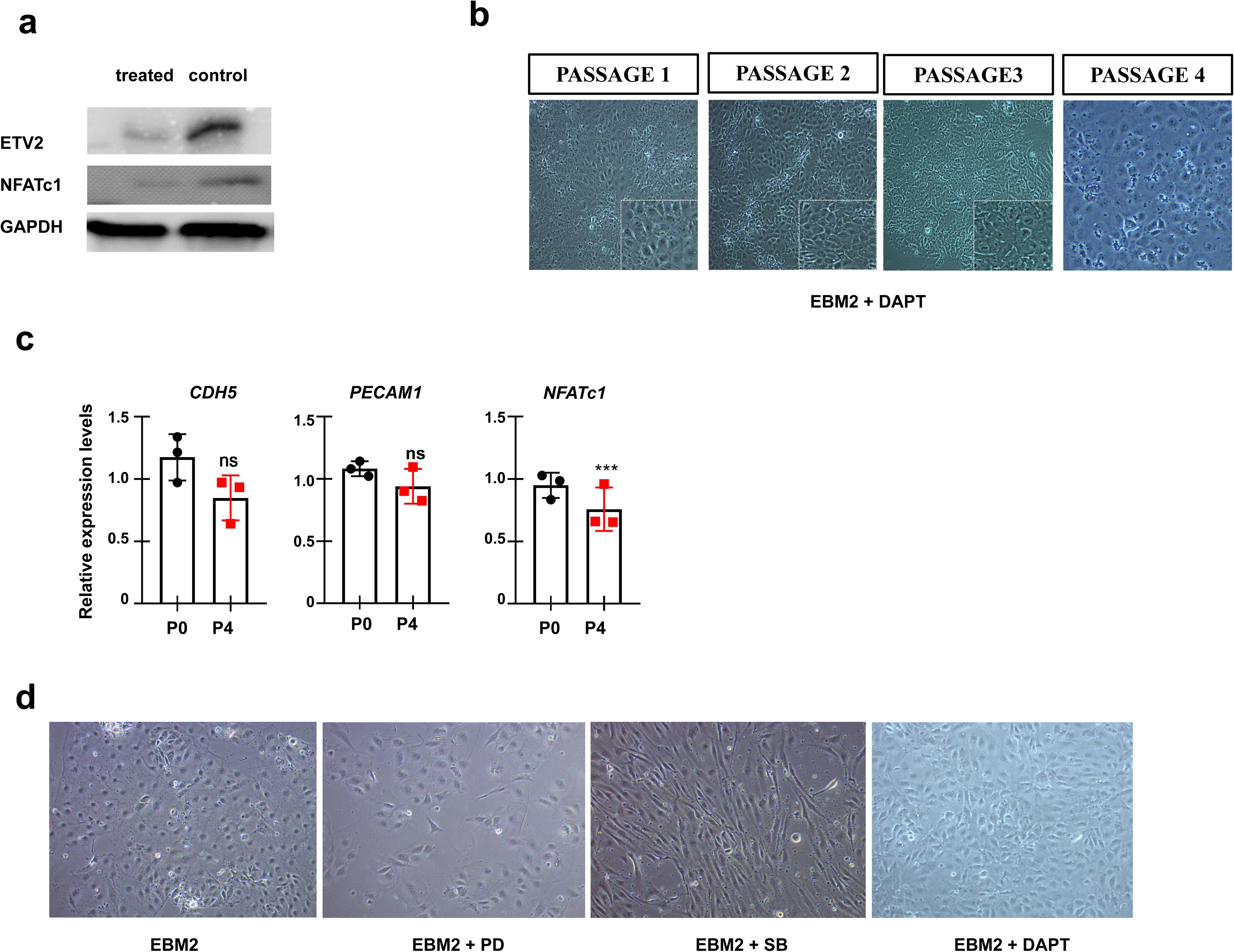
Related to Figure 5. **a** WB analysis of the indicated antibodies for hPSC-derived VEPs treated with and without LDN/SB during differentiation. **b** Morphology of hPSC-derived VEPs of the indicated passage times under the medium EBM2 and DAPT. **c** qPCR analysis of the indicated VEC markers for day 9 hPSC-derived VEPs and hPSC-derived VEPs that have been passaged for 3 times (in EBM2 medium supplemented with DAPT). **d** The morphology of day 9 hPSC-derived VEPs that were cultured for 5-6 days in EBM2 medium in the absence and presence of PD, SB and DAPT. All experiments were repeated 3 times. Significant levels are: \**p*< 0.05; \*\**P*< 0.01; \*\*\**P*< 0.001. NS: not significant. Shown are representative data.

**Supplementary Figure 6.**
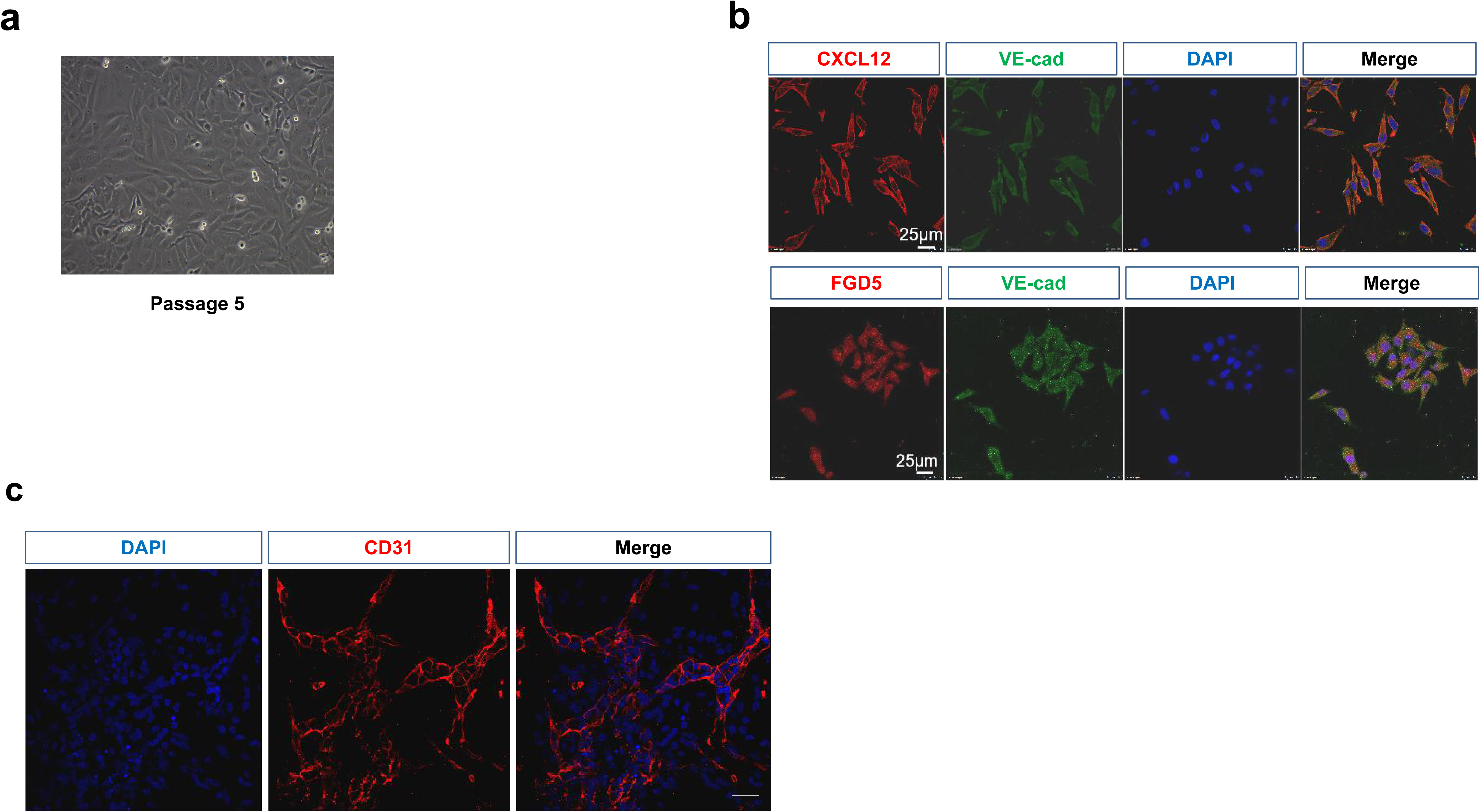
Related to Figure 6. **a** The morphology of hPSC-derived VEPs that have been passaged 5 times. Cells failed to maintain an EC morphology after 5 times passaging. **b** IF staining of the indicated markers for day 9 hPSC-derived VEPs. **c** IF staining of CD31 for day 9 hPSC-derived VEPs. Scale bar: 25 μm. All experiments were repeated 3 times. Shown are representative data.

**Supplementary Figure 7.**
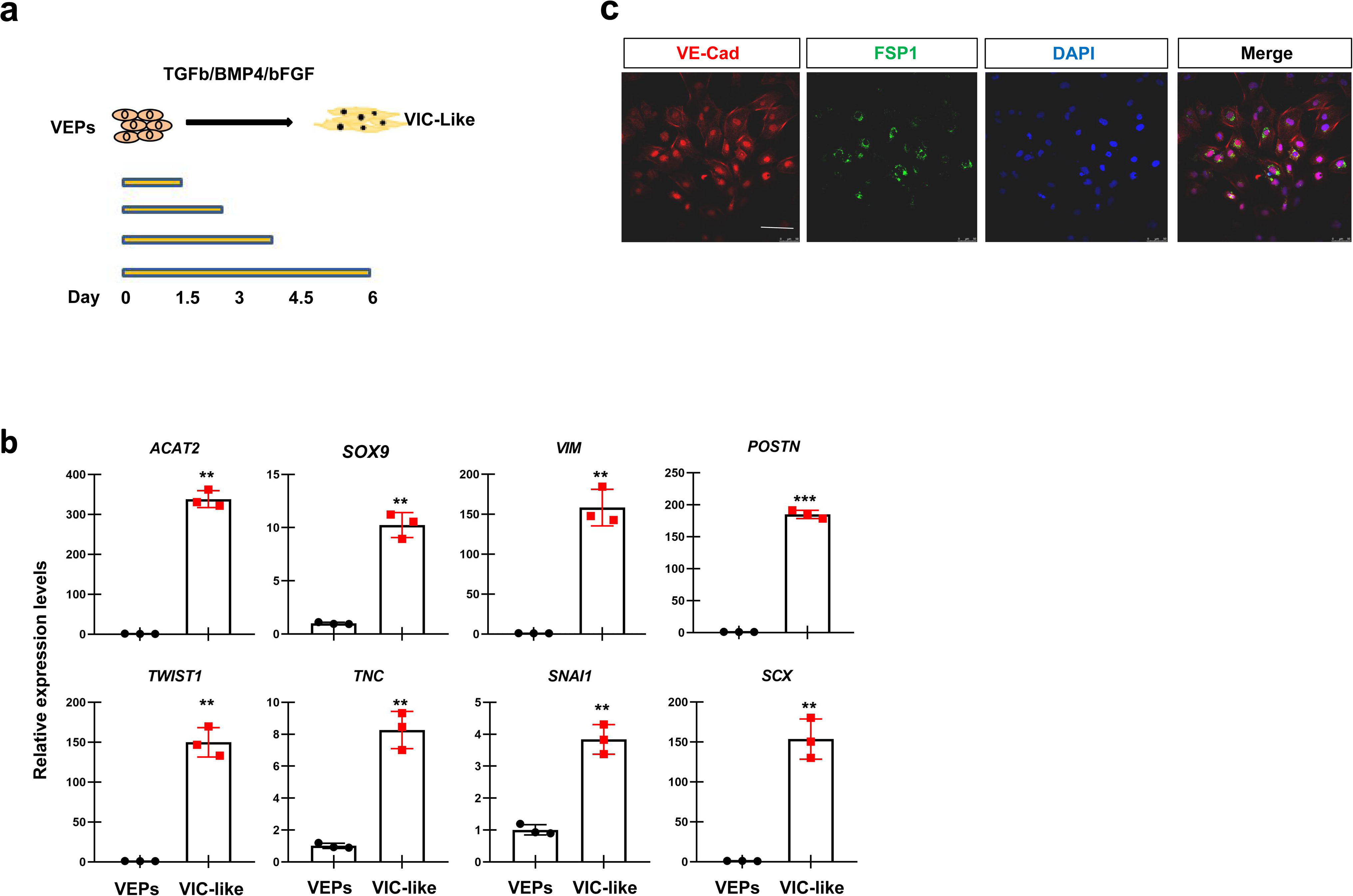
Related to Figure 7. **a** Diagram showing the design of differentiating the VEPs to VIC-like cells. **b** The qPCR analysis of the indicated VIC markers for hPSC-derived VEPs under the combined treatment with TGFb and bFGF for 6 days. **c** IF staining of VE-cad and FSP1 of hPSC-derived VEPs under the combined treatment with TGFb and bFGF for 3 days. Scale bar: 100 μm. All experiments were repeated 3 times. Significant levels are: \**p*< 0.05; \*\**P*< 0.01; \*\*\**P*< 0.001. Shown are representative data.

**Supplementary Figure 8.**
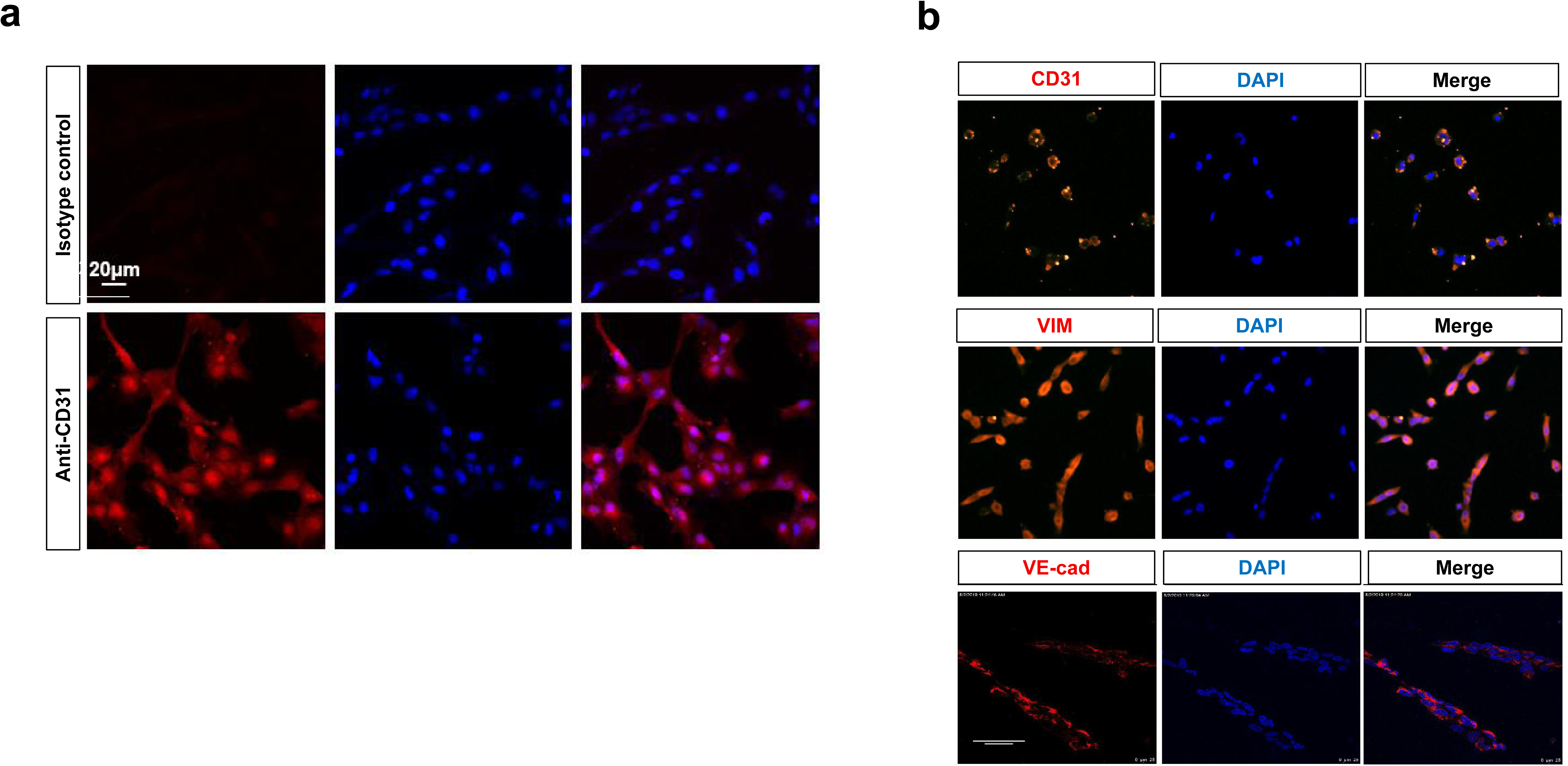
Related to Figure 8. **a** IF signals showing CD31 in hPSC-derived VEPs seeded on the DCVs. **b** IF staining of hPSC-derived VEPs seeded on the DCVs showing the expression of CD31, VE-cad and VIM. Scale bar: 20 μm. All experiments were repeated 3 times. Shown are representative data.

